# Golden pompano genome resource enables discovery of valuable gene determining growth traits

**DOI:** 10.1101/2021.05.20.445047

**Authors:** Honglin Luo, Yongde Zhang, Changmian Ji, Yongzhen Zhao, Jinxia Peng, Yuhui Xu, Xiuli Chen, Yin Huang, Qingyun Liu, Pingping He, Pengfei Feng, Chunling Yang, Pinyuan Wei, Haiyan Yu, Hongkun Zheng, Yong Lin, Xiaohan Chen

**Author notes:** These authors contributed equally to this work (Honglin Luo, Yongde Zhang, Changmian Ji, Yongzhen Zhao, Jinxia Peng, Yuhui Xu). Prime correspondence to: Xiaohan Chen, Hongkun zheng and YongLin are also corresponding authors, **Address Correspondence to:** Mr. Xiaohan Chen, and Mr. Yonglin: NO. 8, Qingshan Road, Guangxi Key Laboratory for Aquatic Genetic Breeding and Healthy Aquaculture, Guangxi Institute of Fishery Sciences, Nanning, 530021, China. Mr. Hongkun zheng: Biomarker Technologies, Beijing, 101300, China.

## Abstract

One important goal of fish genetic breeding is to identify valuable loci and genes that can facilitate growth and thereby productivity. Few such loci or genes have been identified in golden pompano (*Trachinotus ovatus*), a species of significant economic value. In this study, we produced a high-quality chromosome-level genome assembly of the golden pompano by *de novo* sequencing and assemblies for 2 parents and 200 F_1_ offspring by genome re-sequencing. We exploited these assemblies to identify loci and genes by QTL mapping, Kompetitive Allele Specific PCR (KASP) genotyping, and haplotype-based regional association analysis based on growth records of a 64 biparental and 147 individuals from a naturally occurring population. At a locus 291kb from BSNP21031, we identified a somatostatin receptor type 1-like (designated as gpsstr1) gene in which the BSNP1369 of the promoter region was highly associated with growth. Loss of sstr1a, the homolog of gpsstr1 in zebrafish, caused growth retardation. Sstr1a mediated growth via sstr2 and Wnt-gsk-3β signaling pathways. Our findings provide new insights into the underlying mechanisms controlling growth. Our strategy can serve as an effective way to uncover novel genomic information and facilitate improvement of fish growth.

## Introduction

Genetic breeding for higher growth rate, enhanced stress resistance, and better meat quality is becoming indispensable for large-scale commercial aquaculture. Molecular breeding is one of the most promising methods for breeding economically important traits in fish. As one of the most important traits, growth not only determines the efficiency of fish production, but it is also the basis for optimizing the breeding cycle. An improvement in growth rate shortens the cycle, leading to lower costs and higher production. Growth traits are governed by quantitative trait loci (QTLs)^1^ and are influenced by both environmental factors and multiple genes^2^. Benefiting from high-throughput genome sequencing, marker-assisted selection (MAS) using markers linked to QTLs is much more accurate and efficient in improving growth traits in comparison to the traditional selective breeding techniques^3, 4^. Theoretically, if genes and genetic markers associated with growth traits are identified, genetic variants or genes could be used as tools in marker-assisted breeding. Therefore, based on genomic data, genetic linkage maps and molecular genetic marker techniques have been developed to identify the locations of genes associated with growth traits^1^. To date, QTL mapping has been widely used to facilitate the identification of quantitative traits, helping to determine the locations and numbers of linked markers for target traits. Logically, MAS relies on the precise identification of markers tightly linked to QTLs to assist in phenotypic screening. Therefore, accurate identification of QTLs associated with phenotypic traits is a prerequisite for MAS in growth-aimed breeding. Moreover, a high-quality genetic linkage map facilitates QTL mapping for traits with economic value. During the past decade, QTLs for growth traits have been well studied and documented in various economically important fish species, such as common carp (*Cyprinus carpio*)^5^, bighead carp^6^, and Asian seabass (*Lates calcarifer*)^7^. Notably, studies have revealed that growth associated QTLs are distributed on multiple linkage groups in most cases. For example, in Yangtze River common carp, 21 QTLs were distributed among 8 linkage groups^3^, and in *Lates calcarifer*, 6 QTLs in 6 linkage groups were discovered^7^. However, the usefulness of linkage maps regarding the fine-mapping of QTLs has been limited due to their low marker density. Single nucleotide polymorphisms (SNPs) solve such an issue because they can be genotyped on a much larger scale, leading to higher marker density and resolution in the construction of genetic maps^8^. Although genotyping-by-sequencing (GBS) techniques such as 2b-RAD^9^ and SLAF^10^ have been widely applied in discovery of SNP markers throughout the genome^11^, several flaws have been identified in comparison to whole-genome resequencing methods. Whole-genome resequencing produces more SNPs with broader coverage of the genome, thereby facilitating the construction of a higher density linkage map with higher resolution of gene positioning. Therefore, before the emergence of more advanced methods, whole-genome resequencing appears to be a convenient option for QTLs identification and gene positioning.

Golden pompano (*Trachinotus ovatus*, Linnaeus 1758) is a member of the family Carangidae (Rafinesque, 1815)^12^. The species is widely distributed and cultured in the Asia-Pacific region, and it has significant economic importance for offshore cage aquaculture in China and Southeast Asian countries^12^. Due to its high protein content, low fat, and delicious meat, it is being increasingly recognized by consumers worldwide. Golden pompano could be one of the most promising marine culture fish in the future. Notwithstanding the important economic significance of golden pompano, basic research including molecular breeding, immunology, evolutionary biology, and molecular-genetic association studies in this species remain largely unexplored due to the lack of high-quality genomic data and accurate gene annotations. Fortunately, some molecular markers have recently been developed^13, 14^. However, there is only limited information on important economic traits, such as growth, disease resistance, and reproduction, owing to the shortage of genetic information in golden pompano. Moreover, although QTLs identification and gene positioning based on genome sequencing and resequencing have been successfully applied to various plants and animals, few QTLs or genes associated with growth and other traits have been successfully identified in golden pompano. This greatly restricts the process of molecular breeding and the sustained development of the golden pompano industry.

In this study, we performed *de novo* genome sequencing of an adult golden pompano and re-sequenced 202 accessions comprising 2 parents and 200 F_1_ offspring. We developed a strategy to identify loci and candidate genes, and finally identified two SNPs, BSNP21031 and BSNP21028, and one candidate gene annotated as somatostatin receptor type 1-like (designated as gpsstr1) that were significantly correlated with growth. We investigated the functions of gpsstr1 in a zebrafish model. Loss of sstr1a, the homolog of gpsstr1 in zebrafish, caused growth retardation and induced phenotypes that lack the Wnt-gsk-3β signaling. We demonstrated that sstr1a mediates growth via sstr2 and the Wnt-gsk-3β signaling pathway, and the GH-IGF1 axis may regulate the growth defects induced by sstr1a via negative feedback. Our findings provide new insights into the underlying mechanisms controlling growth, and the results will facilitate molecular breeding of fish.

## Results

### *De novo* genome assembly, annotation, and evolution

We first estimated the golden pompano genome size and GC content of golden pompano at 656.98 Mb and 41.54%, respectively, based on 21-Kmer (Figure S1). We then sequenced and assembled the genome of a female individual through a combination of three technologies: paired-end sequencing with the Illumina HiSeq platform, single-molecule real time (SMRT) sequencing with the PacBio Sequel platform, and optical genome mapping with the BioNano Genomics Saphyr System (Figure S2; TableS1, S2A). An assembly containing 1,490 scaffolds with a scaffold N50 length of 21.02 Mb was built (Table S1C and S2A). Next, we anchored and oriented the assembly sequences onto 24 pseudo-molecules, which accounted for ~98.60% (636.61 Mb) of the genome assembly (Figure S3; Table S1 and S2) according to the interaction frequency mapping of 38.7-fold high-through chromosome conformation capture (Hi-C) data (Table S1d). We finally obtained a genome assembly containing 645.62 Mb of genomic data composed of 1,536 scaffolds (including 24 pseudochromosomes) with a scaffold N50 size of 20.22 Mb (Table S2b). The quality evaluation of the assembled genome revealed a high level of contiguity and connectivity for the golden pompano genome and facilitated further analyses (Table S2, S3; Figure S3). By combining the data from *de novo* assembly, homologs, and transcriptome approaches, we annotated 24,186 high confidence protein-coding genes, 152 pseudogenes, and 1,274 noncoding RNAs (Table S3, S4 S5 and S6) in comparison to zebrafish (25,642 protein-coding genes). We visualized the genomic landscape of genes, repetitive sequences, genome map markers, Hi-C data, and GC content of the golden pompano genome using Circos^15^ (Figure 1). As expected, the repetitive elements accumulated in low gene density regions (Figure 1AB). The optical markers and Hi-C data were evenly distributed across the genome (Figure 1CD), as was GC content (Figure 1E). Detailed genomic data can be found in the supplementary information (Table S7-S17).

**Fig. 1.**
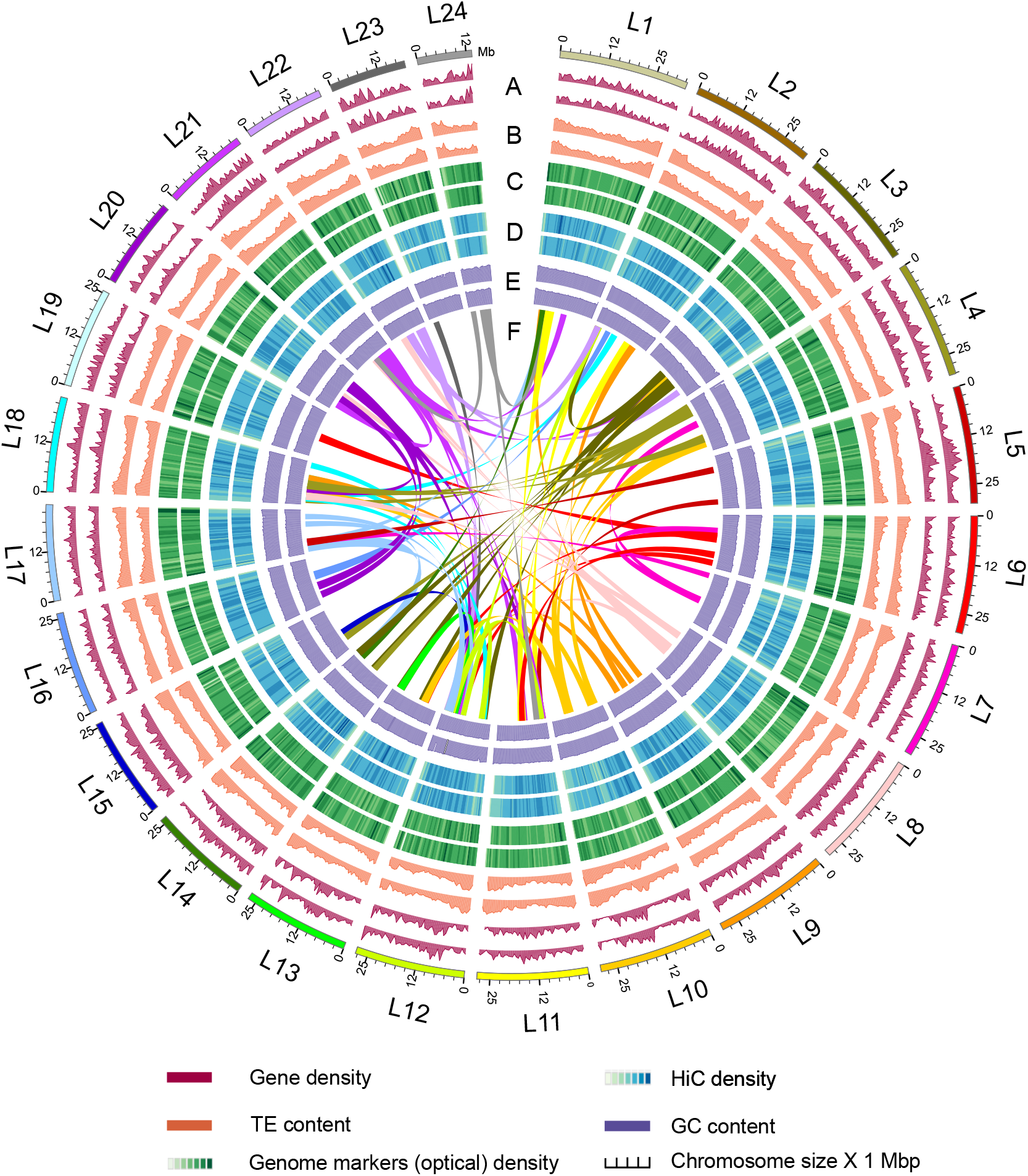
Overview of golden pompano genome (*Trachinotus ovatus*). Numbers on the circumference are at the megabase scale. **A.** Gene density of female *T. ovatus* (window size = 500 Kb). **B.** TE content density of female *T. ovatus* (window size = 500 Kb). **C.** Genome markers (optical) density of female *T. ovatus* (window size = 500 Kb). **D.** Hi-C depth of female *T. ovatus* (window size = 500 Kb). E GC content of female *T. ovatus* (window size = 500 Kb). **F.** Color bands in the middle of the Circos plot connect segmental duplication (minimum five gene pairs) from Teleost-specific whole genome duplication (Ts3R) events.

To examine the evolutionary position of the golden pompano, we reconstructed the evolutionary history of teleost fishes. Phylogenies strongly supported the traditional classification of the golden pompano as belonging to the Carangidae. Golden pompano is a sister of *Seriola dumerilie*, and they diverged approximately 59.99 Mya (Figure 2A). Gene family analyses showed that Perciformes and zebrafish shared 9,412 families, and the number of species-specific families in Perciformes was apparently lower than in zebrafish (Figure S4). Assuming a constant rate of silent substitutions (dS)^16^ of 1.5e-8, we estimated the dates of Ts3R and Ss4R at 350 Mya and 96 Mya, respectively (Figure 2B). Genome collinearity comparison among spotted gar karyotypes (preserved in the golden pompano genome) showed that 894 pairs of paralogous genes that were inherited from the Ts3R event (ohnologues) retained a double-conserved synteny block in the golden pompano genome (Figure 2C, and Table S18), implying that teleost-ancestral karyotypes were considerably conserved in post-Ts3R rediploidization with large fissions, fusions, or translocations (Figure 2D). Next, we classified the Ts3R subgenomes according to the integrity of gene as belonging to the LF, MF, and Other subgenomes^17^. The component of rediploidization-driven subgenomes (LF, MF, and Other) was unequally distributed among golden pompano subgenomes (Figure 2E, and Figure S5), suggesting an asymmetric retention of ancestral subgenomes in teleosts, a phenomenon that is commonly observed in plants^18, 19^. Detail descriptions concerning subgenomes and evolution can be found in the supplementary information.

**Fig. 2.**
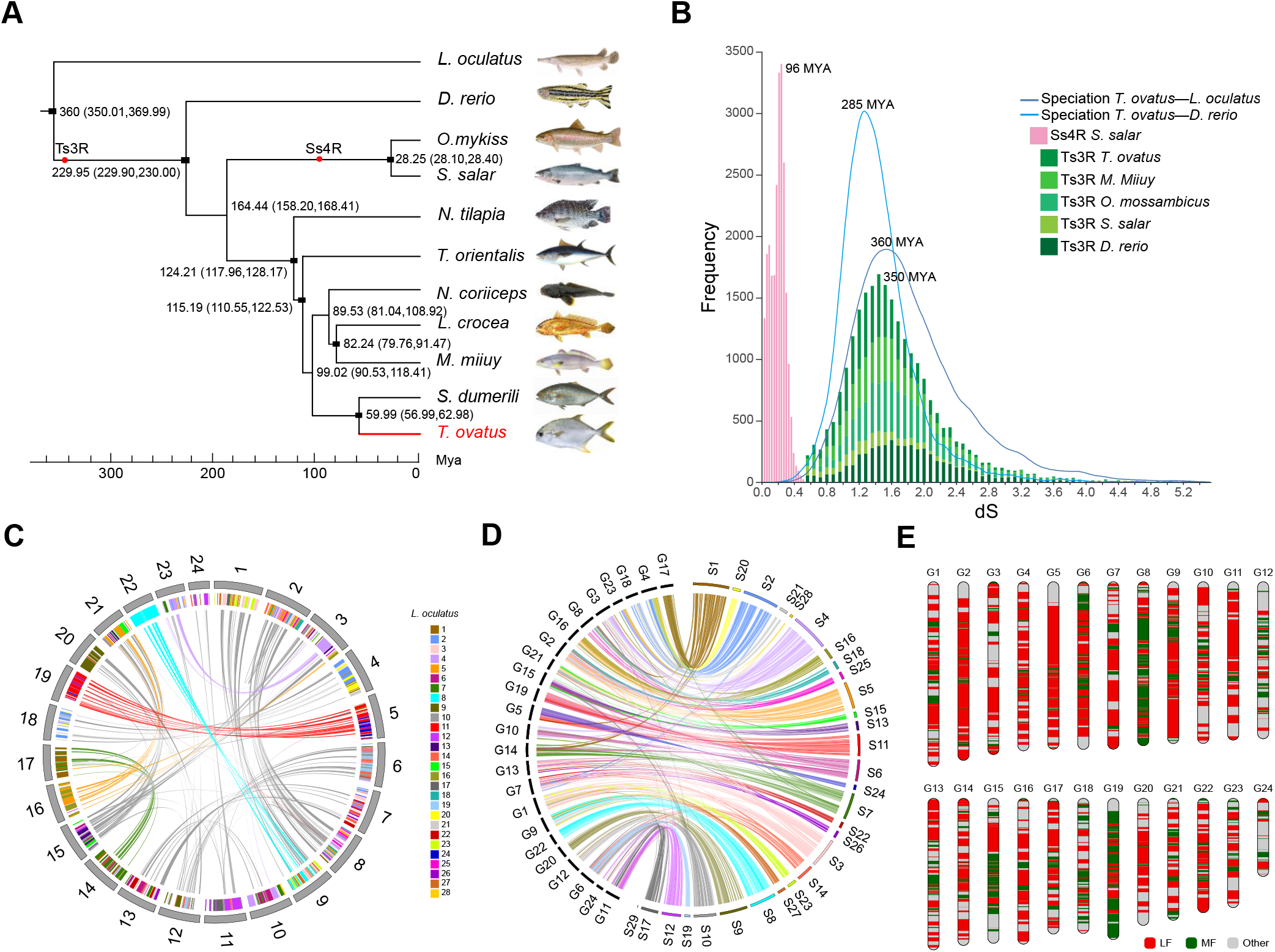
Genome evolution of golden pompano. **A** Phylogenetic relationship of Perciformes and relevant teleost lineages. The position of golden pompano is highlighted in red. Red circles represent the Teleost specific whole genome duplication (Ts3R), Salmonid-specific whole genome duplication (Ss4R), respectively. The divergence time was estimated using the nodes with calibration times derived from the Time Tree database, which were marked by a black rectangle. All estimated divergence times are shown with 95% confidence intervals in brackets. **B** Inspection of whole genome duplication events based on synonymous mutation rate (Ks) distribution. The x axis shows the synonymous distance until a Ks cut-off of 5.2. Note that in order to represent all the data on the same frequency scale, bin sizes are different for each data set. **C** Internal genome synteny of golden pompano. Double-conserved synteny between the golden pompano and spot gar genomes. Each spot gar chromosome (represented as colored blocks) is mostly syntenic with two different chromosomes in the golden pompano genome (syntenic golden pompano regions represented by different colors according to spot gar chromosomal location), a pattern typically associated with whole-genome duplication (Ts3R). Pairs of paralogous genes in spotted gar that are inserted in a double-conserved synteny block are consistent with an origin at the Ts3R event (ohnologues), while genes that are inserted in a double-conserved synteny block but have no paralogue are singletons that have lost their duplicate copy since the Ts3R event. Only genes anchored to chromosomes are represented. **D** Macro-synteny comparison between spotted gar and golden pompano shows the overall one-to-two double-conserved synteny relationship between spotted gar to a post-Ts3R teleost genome. **E** Component of less fragment (LF) and major fragment (MF) subgenomes within golden pompano genome.

### Phenotype variation analysis

We constructed a full-sib family population (FSF), and 200 offspring were randomly selected. Their body weight, full length, body length, and body height were used as indicators of growth traits of the golden pompano, and these phenotypes were identified and recorded at three different time points during cultivation. Among all of the phenotypes, the variance of body weight at the three time points was larger than for other phenotypes, and body weight1026 had the largest kurtosis and CV (Table S19). Comparing the phenotypic data, the other three phenotypes are more stable (Figure 3A; Table S19). Normality analysis showed that Body weight1026, Full length 1011, Full length 1026, Body length 0925 and Body length 1026 were in accordance with normal distributions (Figure 3A; Table S19).

**Fig. 3.**
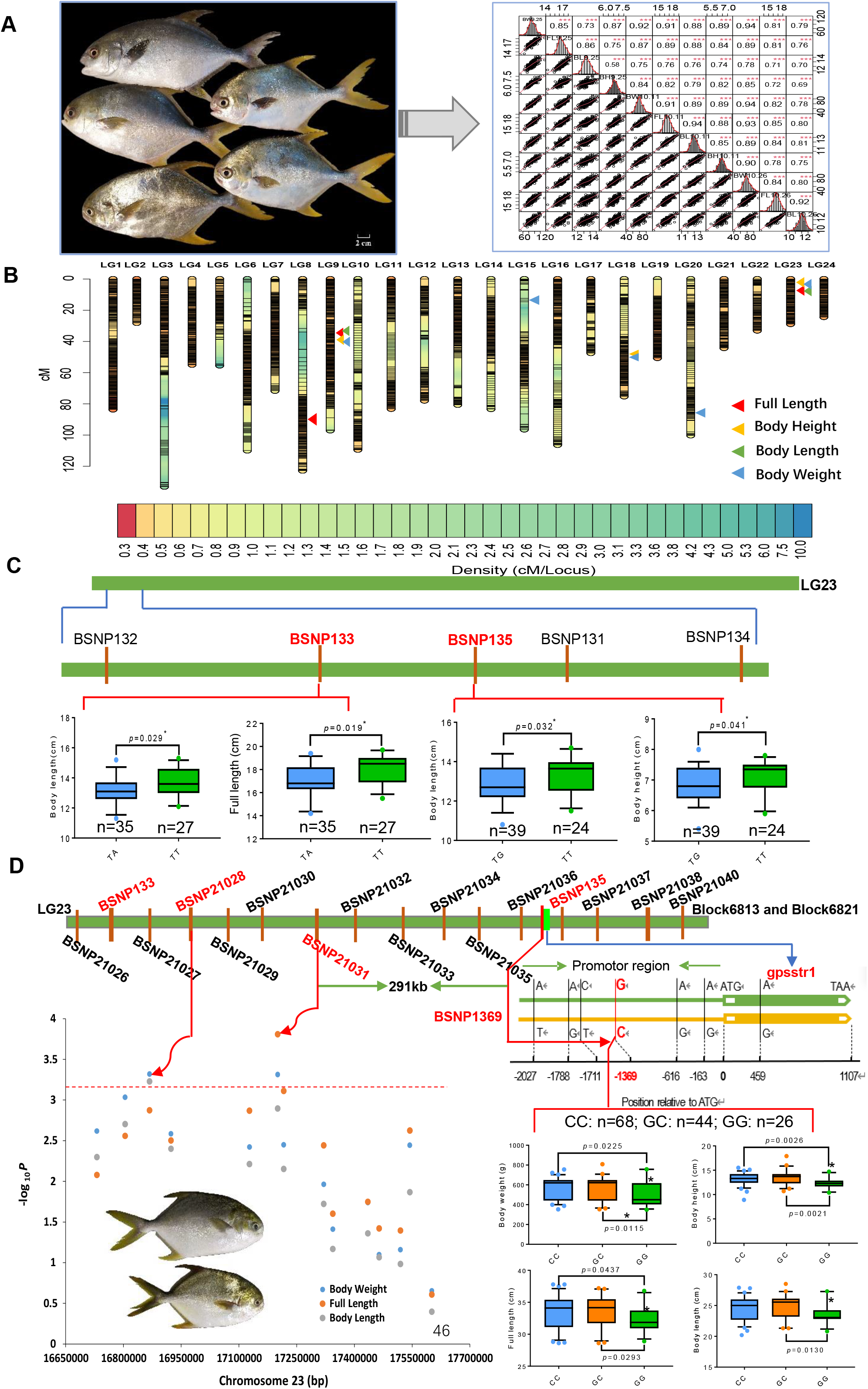
The phenotypes and primary mapping and fine mapping of QTL for growth-related traits of golden pompano. **A.** The appearance of representative F_1_ individuals. Scale bars, 2 cm (down right corner). The variation and Pearson pairwise correlation analyses of body weight (BW), Full length (FL), body length (BL) and body height (BH) of the F_1_ population. The four traits were investigated for three times on September 25^th^, October 11^th^, and October 26^th^ of year 2017, respectively. The correlations were calculated using Spearman correlation coefficients, and the *P* values are indicated as the following: *, *p*< 0.05; **,*p*< 0.01; ***,*p*< 0.001. The analysis was performed using the R package PerformanceAnalytics. Frequency distribution histograms for the four traits are displayed along the diagonal (top right corner). **B.** The high-density genetic map based on 4103 bin markers and repeatable QTLs responsible for BW, FL, BL and BH. The graduated color indicates the marker density on the linkage groups. The triangle of different colors represents different QTLs for corresponding traits. **C.** KASP-based (Kompetitive Allele-Specific PCR) SNP confirmation for QTLs on linkage group (LG) 23. Five evenly distributed SNPs were developed among the cluster QTL region on LG23 and 64 additional F_1_ individuals were genotyped by KASP Assays. The F_1_ individuals were grouped according to their genotypes (TA&TT, TG&TT), and the results were displayed by box charts. The significance testing was conducted by Wilcox Test. **D.** Fine mapping of candidate gene *gpssrt1*. According to the confirmed SNPs (BSNP132, BSNP135), underlying bin marker of Block6813 and Block6821, a total of 14 SNPs were developed to genotype a panel of 146 natural-collected individuals by KASP Assays. Manhattan plot of haplotype-based regional association analysis was displayed. The red dotted line represents the significance threshold (*p*=0.01/14) (bottom left corner). The SNP (−1369) in the promoter region of *gpssrt1* was verified by Sanger sequencing. The natural growing group were grouped according to their genotypes (CC, GC&GG). Two-tailed Student’s *t*-test was used to calculate the *P* value between different groups (bottom right corner). For all the box charts: minimum value = lower whisker, maximum value = upper whisker, median = middle value of box, lower quartile =median of lower half of dataset, upper quartile = median of upper half of dataset, datapoint outside of whiskers = potential outlier.

### Variation detection and genetic mapping

By Illumina re-sequencing, we obtained a total of about 222 Gb clean data. The average sequencing depths of the parents were 14X and 13X, and the average sequencing depth of each offspring was 2X. A total of 579360 SNPs and 171808 InDels were detected between the parents (Figure S6A). Genome annotations showed that more than 40% of SNPs and InDels were located in the intergenic regions. The proportion of synonymous and non-synonymous SNPs in the gene region was 1:1 (Figure S6B), and most InDels caused frameshift mutations (Figure S6C). We next developed bin markers based on SNP data, and a total of 4103 high-quality bin markers were obtained after filtering. The largest bin size was 1.86 Mb, and the average bin size was 91.44 Kb. A genetic linkage map of golden pompano covering 1707.41 cM map distance was constructed by using these bin markers, with an average of 171 bin markers per linkage group and an average genetic distance of 0.41cM (Figure 6C; Table S20). The collinearity and heat map analyses showed that the bin mark order and genetic distance calculation of the genetic map were relatively accurate and could be used for downstream QTL mapping and other analysis (Figure S7).

**Fig. 4.**
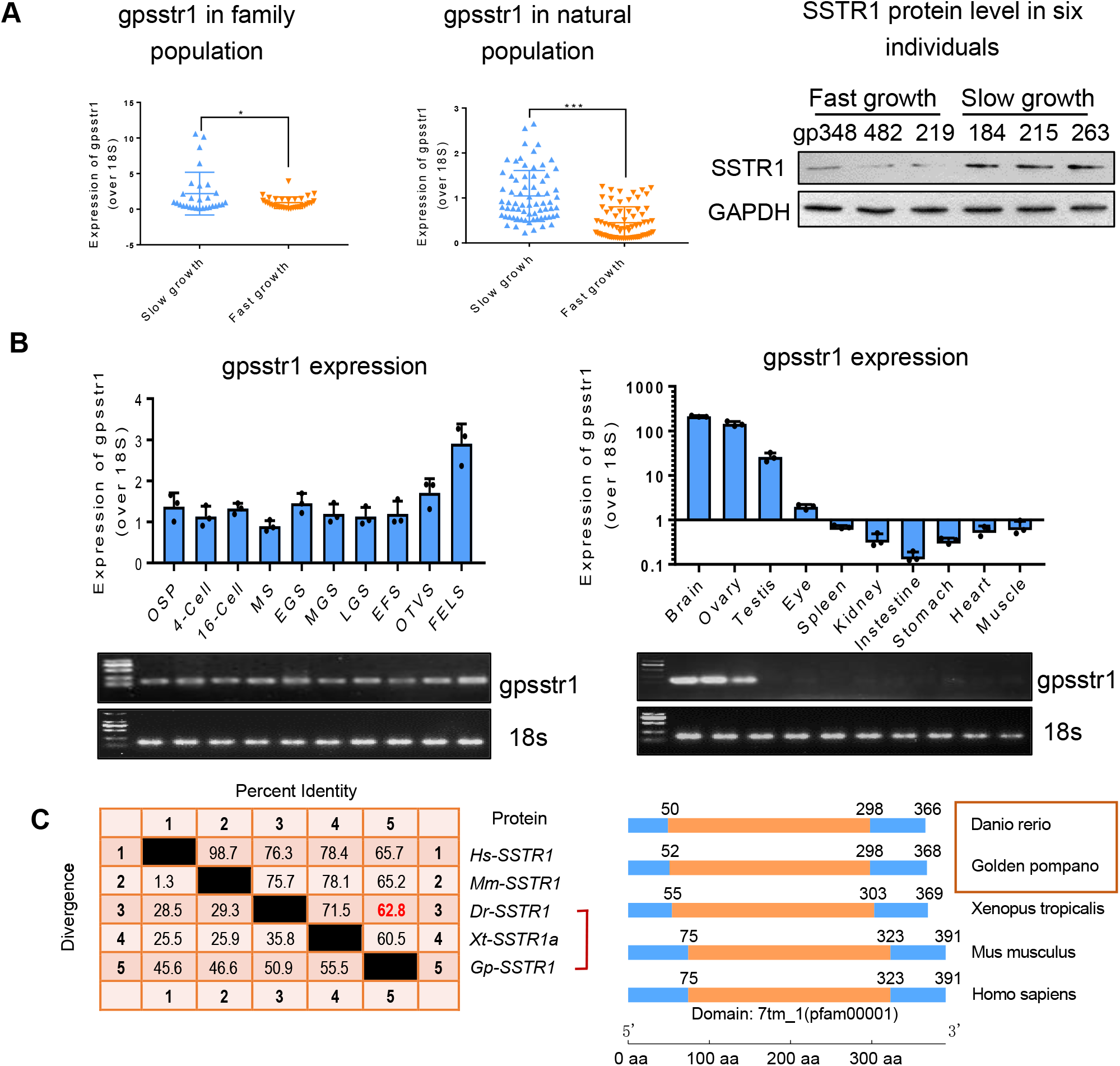
*Gpsstr1*, tightly associated with growth, is a conserved homologue of zebrafish *sstr1a*. **A.** Expression of *gpsstr1* in 64 family population individuals (left) and 146 natural population individuals (middle) of golden pompano, respectively by using qPCR method. Golden pompano fishes were classified into slow and fast growth groups according to their growth trait data. The protein levels of SSTR1 in three fast growth (numbered as gp238, gp482 and gp219) and three slow growth individuals (numbered as gp184, gp215 and gp263) were measured by Western blotting method. Band intensity represents protein level. The data represent as mean±sem. \**p*<0.05 *p*<0.05, \*\**p*< 0.001 represents statistically significant. **B.** Expression profiles of gpsstr1 in 10 embryo stages and 10 tissues of golden pompano by using qPCR (up) and reverse transcription PCR (electrophoretic bands, down). 18S RNA was used as reference. OSP, oosperm; MS, morula stage; EGS, early gastrula stage; MGS, middle gastrula stage; LGS, late gastrula stage; EFS, embryo formed stage; OTVS, otocyst vesicle stage; FELS, formation of eye lens. **C.** *Gpsstr1* is a homologue of zebrafish sstr1a. *SSTR1* protein sequences of *Homo sapien* (*Hs*), *Mus musculus* (*Mm*), *Danio rerio* (*Dr*), *Xenopus tropicalis* (*Xt*) *and Golden pompano* (*Gp*) were aligned by using MegAlign function of Lasergene software (version 6.0). 88.8% identity is found between Gp-SSTR1 and Dr-SSTR1. A conserved domain 7tm_1 was found in both Gp-SSTR1 and Dr-SSTR1 amino sequences.

**Fig. 5.**
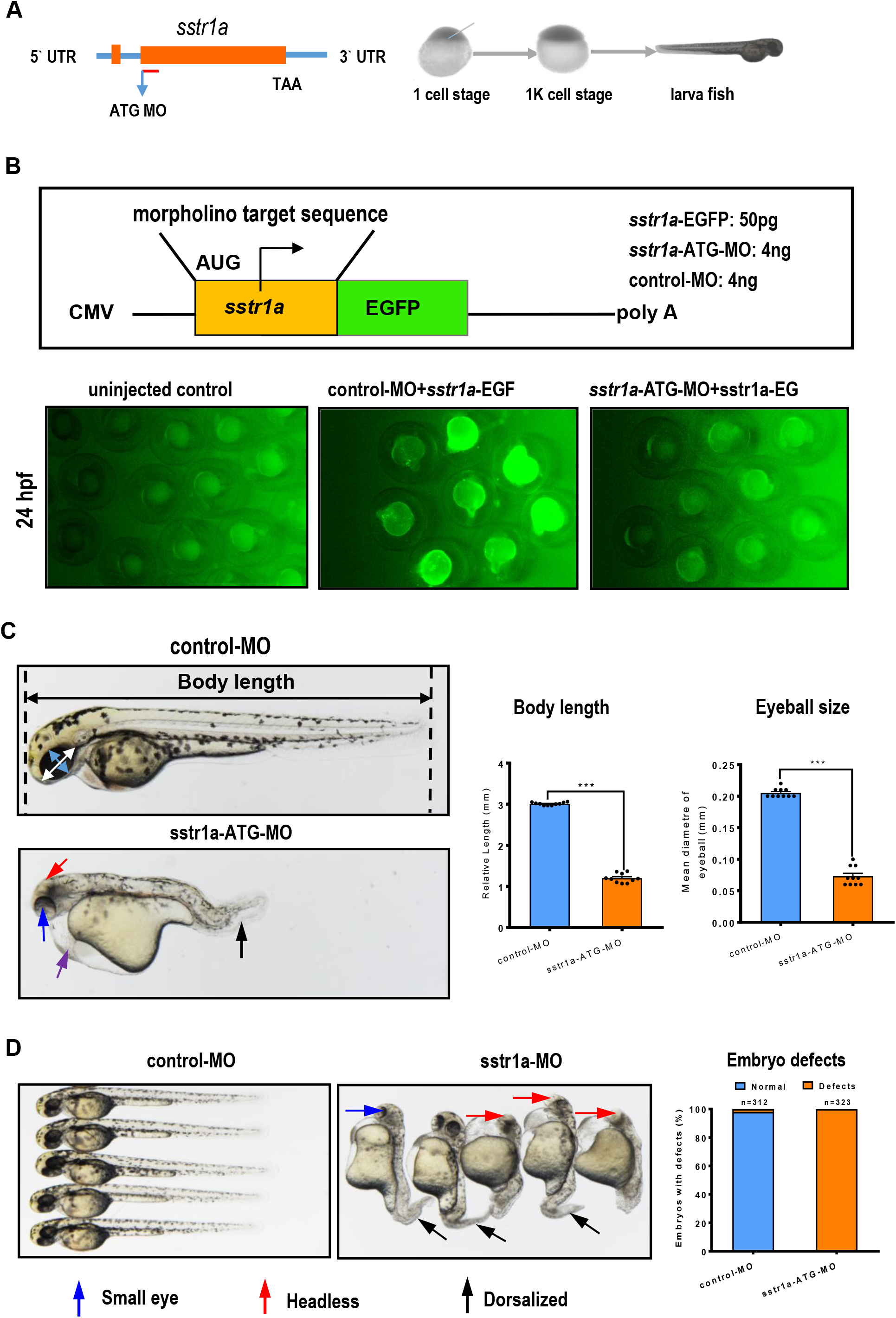
Loss of *sstr1a* delays growth of zebrafish. **A.** Schematic map of *sstr1a* gene and the MO design and micro-injection strategy. **B.** Site-specific effect of *sstr1a-MO* injected into zebrafish. Box shows schematic diagrams of *sstr1a*-EGFP fluorescent reporter mRNAs, the upper one containing the *sstr1a*-ATG-MO target sequence (yellow box) fused in-frame with EGFP. 50pg *sstr1a*-EGFP injected with a standard control morpholino (4ng) or *sstr1a*-ATG-MO (4ng). Embryos were photographed at 24-hpf. Embryos injected with *sstr1a*-GFP plasmid DNA under the driving of CMV promoter showed green fluorescence. When co-injected with *sstr1a*-ATG-MO, green fluorescence decreased dramaticly. hpf, hours post fertilization. **C.** Loss of sstr1a impacts the growth of zebrafish. Body length and eyeball size were measured at 2-dpf. **D.** Gross morphology at 2-dpf. Compared with control MO, knock down *sstr1a* present small eyes (blue arrow), tail patterning defects (dorsalized, black arrow), headless (red arrow) and pericardial edema (purple arrow). The numbers of embryo defects were counted. Error bars, mean ± s.e.m.; \*\*\**p*< 0.0001 (n =10; Student’s t test). dpf, days post fertilization.

**Fig. 6.**
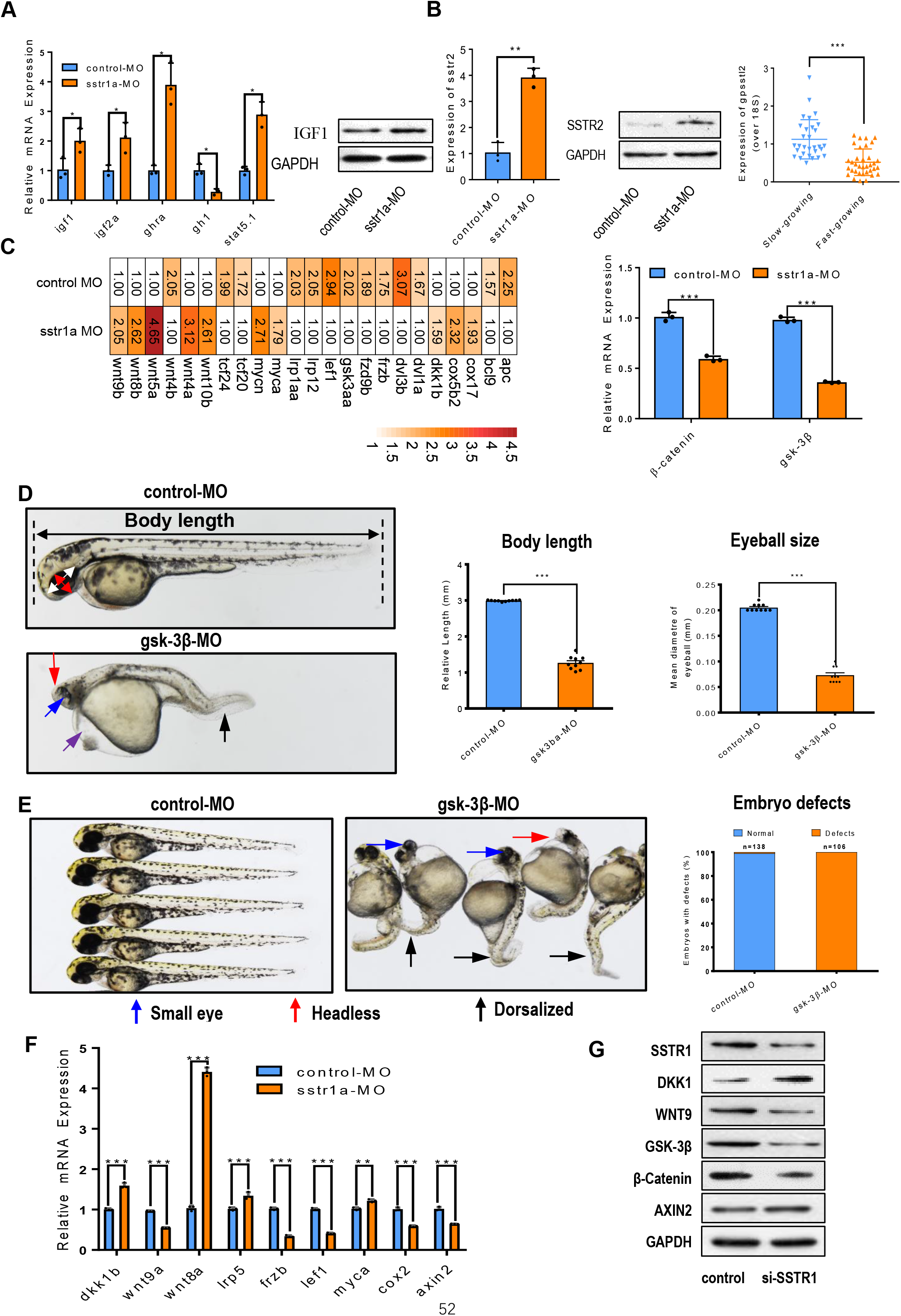
Sstr1 controls growth via Wnt-gsk-3β signaling pathway. **A.** Expressions of growth marker genes in GH-IGF axis. Five genes, igf1, igf2a, ghra, gh1 and stat5.1, were examined by qPCR upon sstr1a depletion in zebrafish. Protein level of IGF1 was also measured by Western blotting with GAPDH as internal reference. **B.** Sstr2 may be responsible to the growth regulation upon sstr1 depletion. mRNA and protein levels of sstr2 were detected by qPCR and Western blotting, respectively. mRNA of golden pompano sstr2 was also measured in 64 family population individuals by qPCR method. **C.** Genes associated with Wnt signaling pathway were enriched upon sstr1a depletion in zebrafish. Heatmap shows the differential expression of each gene in sstr1a knockdown and control group. The bar graph shows the mRNA levels of two key gens, *β*-catenin and gsk-3*β*, in Wnt signaling pathway. **D.** Growth defect phenotype was induced by gsk-3*β* morphant depletion in zebrafish at 2-dpf. Compared with control MO, knock down gsk-3*β* presents small eyes (blue arrow), and decreased both length. The bar graph shows the quantification of the body lengths and eyeball diameters of embryos. These data are representative of three independent experiments. Error bars, mean ± s.e.m.; *** *p*< 0.0001 (n =10; Student’s t test). dpf, days post fertilization. **E.** Gsk-3*β* morphant depletion induced phenotypes resemble those seen in mutant embryos that lack Wnt signaling in zebrafish at 2-dpf and phenocopies the sstr1a depletion. Embryos injected with gsk-3*β* morpholino (MO) displayed typical headless (red arrow) and more anterior truncations (dorsalized, black arrow) than those injected with control MO. These phenotypes resemble the sstr1a depletion. The bar graph shows the percentage of embryos with defects. These data are representative of three independent experiments. **F.** Expressions of key genes involved in Wnt signaling. Nine genes were measured upon sstr1a depletion at 2-dpf by using qPCR method. Compared with control MO, knock down gsk-3*β* significantly downregulated eight genes except the ddk1b and wnt8a. These data are representative of three independent experiments. **G.** Protein expression profiles of genes involved in Wnt signaling. five key proteins were detected upon sstr1 silence in 293T cells by western blotting. GAPHD was used as internal reference. These data are representative of three independent experiments. \**p*<0.05 *p*<0.05, \*\**p* < 0.001 represents statistically significant.

### QTL mapping, fine mapping and candidate gene screening

Next, we wanted to screen a candidate gene that was associated with growth based on the obtained SNPs. We firstly performed QTL mapping by using growth-related phenotypes and genetic linkage maps. A total of 25 QTLs were mapped to the 12 phenotypes and were distributed in linkage groups 6, 8, 9, 14, 15, 16, 18, 20, and 23 (Table S21). Thirteen QTLs could be repeatedly detected, namely, qFL8, qFL9, qFL23 for full length, qBH9, qBH18, and qBH23 for body height, qBL9 and qBL23 for body length, and qBW9, qBW15, qBW18, qBW20, and qBW23 for body weight. There were three overlapping QTL intervals, qFL9, qBH9, qBL9, and qBW9 in linkage group 9, qBH18 and qBW18 in linkage group18, and qFL23, qBH23, qBL23, and qBW23 in linkage group 23 (Figure 3B), implying that there is a phenomenon of one cause for multiple effects in the causal genes of the three intervals. The above three intervals had high confidence, indicating that were the key QTLs that we needed to clone. In order to further confirm and fine map these QTLs, we designed a total of nine KASP markers in these three QTL intervals, including two QTL intervals on linkage group 18, two QTL intervals on linkage group 9, and five QTL intervals in the linkage group 23 (Table S22). We then continued the genotyping in 64 F1 populations and found that the results of these 10 KASP genotyping were consistent with the SNPs results of parental sequencing, indicating the reliability of the SNP sequencing results. However, Wilcoxon tests showed that only BSNP133 and BSNP135 in linkage group 23 were significantly correlated with body length (0.019 ≤ *p* ≤ 0.041, Figure 3C). BSNP133, with a physical distance of 322.17 Kb, and BSNP135, with a physical distance of 351.41 Kb, were located in Block 6813 and Block 6821, respectively. In order to further confirm this positioning, we collected 147 individuals from a natural population and designed 14 evenly distributed KASP markers between Block 6813 and Block 6821 (between 16730073-17600034 on the genome). We found that the peak SNP BSNP21031 (17199635) located in Block 6821 was extremely significantly correlated with body weight and full length, and BSNP21028 (16866882) located in Block 6813 was extremely significantly correlated with body weight and body length (Figure 3D) by using hapQTL association analysis. No non-synonymous mutations were found in the Block 6813 interval between the parents. In Block 6821, only one non-synonymous mutation site on EVM0019027 was found, but no GO or KEGG terms were enriched. At a distance of 291kb from the BSNP21031 locus, we found a gene EVM0001630 that was annotated as somatostatin receptor type 1-like (K04217) with somatostatin receptor activity (GO:0004994) in Block6821, which clearly suggests a role in the growth process (Table S23). We further found seven SNPs near gene EVM0001630 (17491064-17492170) among which there was a mutation BSNP1369 (G→C, 17489695, Figure 3D) in the promoter region of EVM0001630. This SNP was verified by Sanger sequencing of the promoter region of EVM0001630 (Table S24) and was significantly correlated with body weight, body height, full length, and body length (Figure 3D). This indicates that the gene EVM0001630 with SNP (BSNP1369) in the promotor may be a candidate gene that is associated with growth.

### *Gpsstr1*, tightly associated with growth, is a conserved homologue of zebrafish *sstr1a*

We designated EVM0001630 (somatostatin receptor type 1-like) gene as gpsstr1. Then, we asked whether gpsstr1 was associated with growth of golden pompano. First, we detected the mRNA level of gpsstr1 in 64 individuals and found that it was significantly higher in fish growing slowly than those growing rapidly. Similarly, its expression level was much higher in slow-growing individuals than fast-growing fish in individuals from a natural population. We next examined the protein level of SSTR1 in six fast- and slow-growing fish selected from the 64 family groups by Western blotting, and the protein expression profiles resembled the mRNA profiles in the family or natural populations (Figure 4A). These data imply that gpsstr1 is tightly associated with growth of the golden pompano. We also detected the distribution of gpsstr1 in golden pompano by qPCR and RT-PCR, and we found that it was widely distributed in all of the tested embryo stages, with much higher expression in the formation of the eye lens (FELS) embryo stage, where it was highly expressed in brain, ovary (females), and testis (male) but not in the eye, spleen, or heart (Figure 4B). We speculated as to its biological functions; this needed an ideal fish as a research model. The zebrafish has long been used as such an ideal model, so we needed to address whether there was a homolog of gpsstr1 in the zebrafish. We searched for a homolog of gpsstr1 in the NCBI database by BLASTp: only one gene, sstr1a (mRNA: XM_691574.8; protein: XP_696666.1) shared 62.8% similarity at the amino acid level in zebrafish (Figure 4C). Both proteins possess the same conserved domain 7tm_1 (pfam0001) and SSTR1 protein signature motifs YANSCANPILY and DRY (Figure S8). Also, the simulated 3D structures of both proteins were highly similar (Figure S9). These results suggested that gpsstr1 and zebrafish sstr1a are highly conserved and that they may have similar biological functions. Therefore, we speculated as to the functions of sstr1.

### Loss of sstr1a causes growth defects in zebrafish

Next, we addressed the function of sstr1 in fish growth by using the zebrafish model, a system that has been a valuable tool for functional research in vertebrates due to its high similarity in fishes and other vertebrates^20^. First, we investigated whether loss of *sstr1a* affects growth traits in zebrafish using a morpholino micro-injection strategy (Figure 5A). We constructed a pcDNA3.1-sstr1a-EGFP fluorescein reporter system containing sstr1a-ATG-MO (morpholino, MO; sstr1a contains only one exon, and thus only ATG-MO was designed) and tested its site-specific effect when injected into zebrafish at 24 hours post fertilization (hpf). As shown in Figure 5BC, compared to the control embryos, the green fluorescence in embryos that were co-injected with *sstr1a*-ATG-MO decreased dramatically, confirming the high efficiency of sstr1a-ATG-MO (Figure 5BC). Next, we examined the effects of sstr1a depletion on the growth of zebrafish, and we observed that sstr1a depletion caused growth defects manifested as significant decreases of body length and eyeball size compared to controls (Figure 5D). Furthermore, loss of sstr1a resulted in a high percentage of embryo defects (100%, n=323) and caused phenotypes such as small eye, dorsalized, headless, and pericardial edema at two days post fertilization (dpf) compared with controls (Figure 5E). This confirmed that knockdown of sstr1a affects growth in zebrafish.

### The GH-IGF axis may negative regulate sstr1-induced growth defects

The growth hormone-insulin-like growth factor (GH-IGF) system plays a major role in coordinating the growth of fish^21, 22^. Numerous research studies of fish growth have focused on the growth hormone (GH), a component of the GH-IGF system of fish. This system not only includes GH receptors (GHR), IGF receptors (IGFR), GH binding proteins (GHBP), and IGF binding protein (IGFBP), but it also consists of multiple subtypes of GHRs, IGFBPs, and IG-FRs^23, 24^. These components are broadly distributed and interact to coordinate growth and other essential biological processes. As the GH-IGF1 axis has been demonstrated to be closely associated with hormone production and affects growth processes in vertebrates^23^, we supposed that the sstr1a-caused growth defects might also be associated with the GH-IGF axis. We next examined the expression levels of the growth markers igf1, igf2a, ghra, gh1, and stat5.1 in zebrafish. We found that sstr1a depletion resulted in a significant decrease of gh1 expression compared with normal fish. However, loss of sstr1a significantly increased the expression levels of igf1, igf2a, ghra, and stat5.1 at the mRNA level and IGF1 at the protein level compared to controls (Figure 6A). RNA-seq data also corroborated this upregulation tendency of some GH-IGF-related genes, including ghrb, igf2a, and stat5a, upon sstr1a depletion, (Table S25). The decrease of sstr1a released the inhibition of the GH-IGF system, and this consequently resulted in larger body length and eyeball size. However, our fish exhibited opposite phenotypes. This may suggest that sstr1a does not directly act via regulation of the GH-IGF system. Given that the GH-IGF axis positively regulates growth^23^, our data demonstrate that this axis more likely negatively regulates the growth phenotypes induced by sstr1a knockdown.

### Sstr1 mediates growth via sstr2 and the Wnt-gsk-3β signaling pathway

Since the GH-IGF could not fully explain the growth phenotypes such as body length decrease and eyeball diminution, we next examined the expression of sstr2, a protein that forms a heterodimer with sstr1^25^ and that is negatively regulated by sstr1^26^ and positively associated with growth regulation^27^. We found that when sstr1a was knocked down in zebrafish, the mRNA expression of sstr2 increased significantly compared with the controls, and the protein level of sstr2 also increased when sstr1a was silenced (Figure 6B). This is consistent with the previous research reporting that loss of sstr1 in mice resulted in an increase of sstr2 retinal expression as well as enhanced sstr2 function. This suggests an inhibitory role of sstr1 on sstr2 expression^26, 28, 29^. As it has been reported that sstr2 plays a more important role in regulating animal growth than sstr1^27^, we wondered whether golden pompano sstr2 (designated as gpsstl2) was associated with growth. The sstr2 mRNA levels in fast- and slow-growing golden pompano were quantified by qPCR; the results showed that it was significantly higher in slow-growing fish than that in fast-growing fish. This strongly suggests that gpsstl2 is tightly related to growth of golden pompano, consistent with previous research^27^. This also suggests that gpsstl2 rather than gpsstr1 may directly control growth in golden pompano. Therefore, the upregulation of zebrafish sstr2 in our study partly explains the retarded growth induced by sstr1a depletion. However, this could not explain the phenotypes such as small eye, dorsalized, and headless in zebrafish. We noticed that these phenotypes are characteristic of dorsalization and resemble those phenotypes seen in embryos that lack Wnt signaling^30, 31^. This suggests that sstr1a might function via the Wnt signaling pathway. To validate this conjecture, we analyzed the RNA-seq of zebrafish embryos upon sstr1a depletion and found that the transcriptome data presented a large amount of differentially expressed Wnt-signaling genes enriched after sstr1a depletion (Figure 6C; Figure S10; Table S25), suggesting that loss of sstr1a affects the Wnt signaling pathway. Next, we examined the key genes in the Wnt-signaling pathway, *β*-catenin and gsk-3*β*, and found that both were decreased significantly upon sstr1a depletion compared with controls (Figure 6C). To further verify our hypothesis, we knocked down the gsk-3*β* gene in zebrafish using the morpholino technique to determine whether loss of gsk-3*β* phenocopies sstr1a deficiency. After testing the efficiency of gsk-3*β*-MO (Figure S11), we injected 4 ng gsk-3*β*-MO in embryos and observed that loss of gsk-3*β* caused growth defects manifested as significant decreases of body length and eyeball size compared to controls (Figure 6D). Furthermore, gsk-3*β* depletion resulted in a high percentage of embryo defects (100%, n=106) and produced the phenotypes small eye, dorsalized, headless, and pericardial edema at 2 dpf (Figure 6E) compared with controls. These results indicate that gsk-3*β* depletion not only led to growth defects similar to those observed from sstr1a depletion (decreased body length and eyeball size) but also phenocopied the sstr1a deficiency (small eye, dorsalized, headless, and pericardial edema). This suggests that sstr1a functions via the Wnt-gsk-3*β* pathway. To further validate this, we examined the mRNA expression of genes in the Wnt-signaling pathway upon sstr1a depletion, and we found that most of the key genes including wnt9a, irp5, frzb, lef1, myca, cox2, and axin2 were significantly downregulated, while dkk1b and wnt8a were sharply upregulated compared to controls (Figure 6F). Finally, we silenced the sstr1 gene in 293T cells using siRNA and monitored the expression levels of the key proteins by Western blotting. The bands showed that the protein levels of WNT9A, GSK-3*β*, *β*-Catenin, and SSTR1 decreased, while DKK1 increased. This was consistent with the corresponding mRNA expression profiles, except for AXIN2 that was almost unchanged at the protein level (Figure 6G). Based on these data together with evidence of heterodimerization of sstr1 and sstr2^25^, we concluded that sstr1a plays a role in growth regulation through sstr2 and the Wnt-gsk-3*β* signaling pathway. To this end, we attempted to delineate the possible mechanism by which sstr1 mediates growth by bridging the Wnt signaling and GH-IGF axis (Figure 7). Briefly, our data indicated that sstr1 depletion may on the one hand activate sstr2 and decrease gh1 to inhibit growth and upregulate igf1, igf2a, and ghra to suppress such over-inhibition via negatively feedback. On the other hand, sstr1 depletion downregulated Wnt-gsk-3*β* signaling, which explains the phenotypes of growth delay, small eyes, pericardial edema, and tail pattering defects observed under sstr1 depletion.

**Fig. 7.**
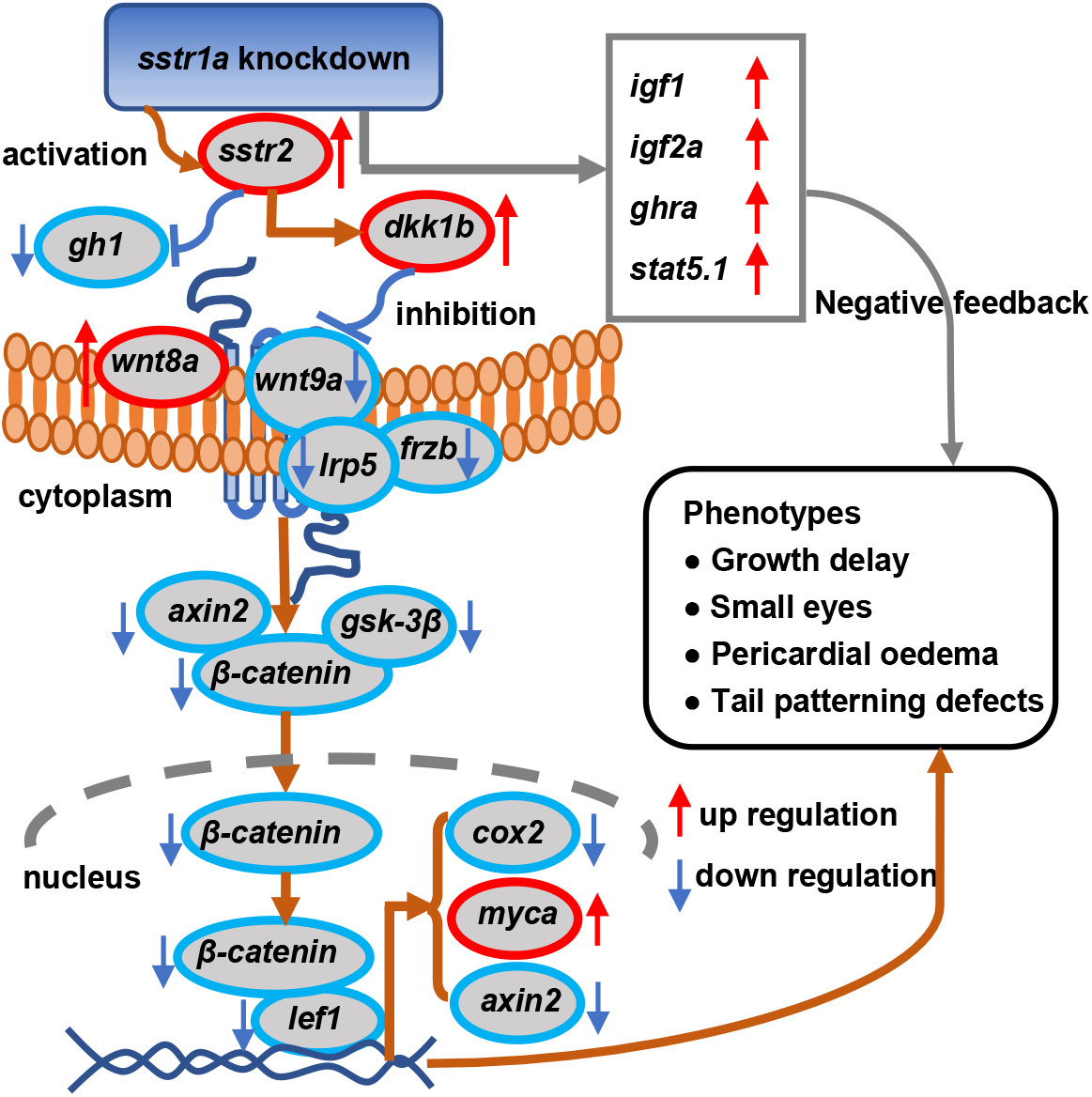
Schematic illustrating the possible mechanism that how sstr1 mediates growth by bridging Wnt signaling and GH-IGF axis. Sstr1 depletion may on one hand activate sstr2 and decrease gh1 to inhibit growth, and on the other hand upregulate igf1, igf2a, ghra to negatively feedback suppress such over-inhibition. On the other hand, sstr1 depletion downregulates 9 key genes playing essential roles in Wnt-gsk-3*β* signaling. The downregulation of Wnt-gsk-3*β* signaling explains the phenotypes of growth delay, small eyes, pericardial edema and tail pattering defects observed by sstr1 depletion.

## Discussion

In this study, we obtained a high-quality chromosome-level genome of the golden pompano by performing *de novo* genome sequencing. This genomic resource not only provides a high-value reference genome for functional genomics studies of golden pompano, such as exploration of economic traits and sex determination, but it also offers an important platform for studies in other fundamental fields such as immunology, molecular biology, and evolutionary biology. We also found that the golden pompano genome, as in other teleosts, underwent the Ts3R whole-genome duplication (WGD) event that resulted in biased-subgenomes retention. This also provides clues for research on fish physiological and morphological evolution^32^, evolutionary embryology, and WGD-derived subgenome evolution^33, 34^. Therefore, from the point of view of basic research, this genomic resource may help to strengthen the development of the golden pompano industry.

One important goal of genomic and genetic studies of cultured fishes is to identify important loci and genes that can be used to improve economic traits and thereby productivity. Growth determines the efficiency of fish production. Growth-related traits are governed by quantitative trait loci (QTLs). In this study, in order to map causal genes responsible for growth-related traits, we designed an experimental process that combined construction of a biparental population and a natural population with whole-genome resequencing-based QTL mapping methods and haplotype-based regional association analysis. The process includes: (i) the detection of QTLs through linkage analysis with genome-wide nucleotide polymorphisms; (ii) the confirmation of QTLs with SNP-derived Kompetitive Allele Specific PCR (KASP) markers; (iii) the QTL fine mapping of significant signals based on association analysis with newly designed KASP markers using the natural population; (iv) the extraction of causal genes based on the annotation of polymorphisms and function enrichment information (GO and KEGG), including that of zebrafish homologs as in the case of EVM0001630 (designated as gpsstr1); (v) candidate gene confirmation through DNA cloning by Sanger sequencing; and (vi) functional and mechanistic analysis of candidate genes in a zebrafish model. As expected, we identified two SNPs in steps ii and iii after detection of 25 QTLs, BSNP21031 and BSNP21028, that were associated with body weight and full length and body weight and body length, respectively. Near BSNP21031(about 291kb), we extracted one causal gene, gpsstr1, that was annotated as somatostatin receptor type 1-like, implying its important role in the growth process. Notably, one SNP, BSNP1369 in the promoter region of gpsstr1, was significantly correlated with growth of golden pompano, supporting the conclusions from the above strategies. To this end, we finally identified a target candidate gene, and this indicated that our QTL mapping and fine mapping strategy using F_1_ individuals together with fish from a natural population was reliable and feasible. To our knowledge, this is the first report of QTL fine mapping and candidate gene screening on growth traits of golden pompano (*T. ovatus*) based on whole-genome sequencing and re-sequencing data, although several genetic markers have been documented previously^35, 36^.

Growth is a polygenic trait influenced by multiple physiological pathways. Genes of the GH-IGF axis are thought to have the greatest influence on growth among the possible growth-regulating pathways in vertebrates (including fishes). A number of growth hormones such as somatostatin, insulin, and sex steroids modulate the growth process in fishes via the GH-IGF system^21^. Recently, polymorphisms of the QTL-related candidate genes associated with the GH-IGF system have been demonstrated to be closely associated with growth in fish, for example GHR2 in tilapia^37^ and GH in Arctic char^38^. As one of the most important inhibitors of the GH-IGF system, somatostatin receptor (sstr) genes have seldom been reported in association with growth-aimed breeding in fish. Sstrs participate in regulation of a variety of hormones. Their regulatory effects are closely related to growth, neurology, immunity, and disease^27^. It is significant to study their functions, mechanisms, and relationships with growth. At present, research on sstrs is mainly focused on mammals, with few studies in fishes. Five sstrs have been identified in mammals. Sstr2 and sstr5 rather than sstr1 and sstr3 play important roles in regulation of animal growth and inhibition of GH and insulin^27^. In fishes, only four sstrs (sstr1, 2, 3, and 5) have been discovered. Studies of their association with growth have seldom been reported, except for a study on the association of sstr2 with body size in African cichlid fish^39^. In this study, we identified the gpsstr1 gene as one of the sstr subtypes of fish not only because it possesses a conserved seven transmembrane domain but also through the signature motifs YANSCANPILY and DRY, as all sstrs possess^27^. Our data showed that gpsstr1 was negatively associated with golden pompano growth in both F_1_ fish and those from a natural population. The gpsstl2 expression exhibited a similar profile to that of gpsstr1. This is consistent with the general characteristics of sstr1 and sstr2 having inhibitory functions on the growth process^40, 41^. To the best of our knowledge, this is the first growth-related phenotype in sstr1-deleted zebrafish, although phenotypes of altered glucose and insulin homeostasis were previously documented in sstr1-ablated mice^42^. However, the phenotypes induced by sstr1a depletion in zebrafish were outside the expected forms given the inhibitory function of sstr1 on growth. The depletion of sstr1 was expected to lead to rapid growth. However, loss of sstr1a caused opposite phenotypes. This implies that additional factors may be involved in this process, or that sstr1a plays different roles during embryo and adult stages. Fortunately, we found that sstr1a depletion activated sstr2 and decreased gh1expression. This is consistent with the previous report that sstr2 activation is related to inhibition of GH release in fish^27, 41, 43, 44^. Also, previous research suggested that sstr2 plays a much more important role in growth regulation than sstr1^27^. Consequently, in this study the activation of sstr2 upon sstr1 depletion enhanced the inhibition of GH1 release; this explains the retarded growth phenotypes. However, loss of sstr1a simultaneously increased the other four key genes in the GH-IGF axis. Their upregulation could not explain the retarded growth phenotypes. This implies the existence of new downstream signaling pathways of sstr2 in addition to the GH-IGF axis. We found that the upregulation of the key genes in the GH-IGF axis was not the main mechanism for the phenotypes induced upon sstr1a depletion. The decreased inhibition of sstr1a is supposed to result in the upregulation of GHs; however, this upregulation should logically promote and not hinder growth. Considering the complexity of growth regulatory networks, based on our current data, we prefer to consider that such upregulation is a negative feedback mechanism in which the sstr1a depletion induced retarded growth. More investigation is needed to clarify the mechanism behind these results; this will be a focus of future work.

Our study highlights the important discovery of regulatory roles of sstr1 on the downstream Wnt-gsk-3*β* signaling pathway. Wnt signaling plays multiple roles in the control of cell proliferation, differentiation, specification, and growth and maturation of most tissues during embryonic development and adult tissue homeostasis in animals^45, 46, 47, 48, 49, 50^. As supporting evidence, a novel role for Wnt and TGFβ in regulating size-dependent growth was recently identified in planarian flatworms, highlighting the function of Wnt signaling on growth modulation by coordinating other signaling pathways^51^. Previous studies showed that the mammalian sstrs are mainly involved in the following pathways: cyclic adenylate (cAMP), voltage-dependent Ca^2+^ channel, mitogen-activated protein kinase (MAPK), and protein tyrosine phosphatase (PTP) pathways^27^. Still, the downstream signaling pathways of sstrs are largely unclear. To date, no direct evidence has been provided for the correlations of sstrs with the Wnt-gsk-3*β* pathway in regulation of animal growth. Only one previous report mentioned that Octreotide (one somatostatin analog peptide) but not sstr activates gsk-3*β*^52^. In our study, the unique phenotype of dorsalization induced by sstr1 depletion resembled the signature phenotype of Wnt signaling^30, 31^. Such a dorsalization phenotype has never been documented in sstr1-ablated mice or other animals. Moreover, loss of gsk-3*β* duplicated the sstr1a-induced phenotypes, together with the significant downregulations of key gens in Wnt signaling pathway, again strongly suggest that sstr1 mediates growth through the Wnt-gsk-3*β* signaling pathway. In our study, although the lack of commercial antibodies of zebrafish and golden pompano impeded the examination of protein levels of sstr1 and key genes in the Wnt-gsk-3*β* signaling pathway *in vivo*, the Western blotting in 293T cells supported our hypothesis that sstr1 mediates Wnt signal transduction. To the best of our knowledge, this is the first report of sstr1 regulating downstream Wnt-gsk-3*β* signaling pathway. From this point of view, our study may open a new window on sstr1 research, although some questions need to be answered in the future, such as whether more genes are involved, which genes in the Wnt pathway directly interact with sstr1, and how the GH-IGF axis cross-talks with the sstr1-Wnt-gsk-3*β* signaling pathway.

In summary, we have developed an efficient strategy that combines construction of a biparental population and a natural population with whole-genome-resequencing-based QTL mapping methods and haplotype-based regional association analysis. This strategy can greatly enhance the efficiency of uncovering QTLs and genes for beneficial economic traits in fish. By using this strategy, we were able to identify a candidate gene for growth regulation, and we validated the candidate via gene knockdown tests in a zebrafish model. Notably, to our best knowledge, this is the first report identifying the candidate gene sstr1 as having a role in the growth process via regulation of downstream Wnt-gsk-3β signaling pathway as well as the first report of headless and dorsalization phenotypes caused by sstr1a. Clearly, this study has provided valuable new insights into the underlying mechanism of the growth process. Our study will further enable the molecular manipulation of desirable economic traits of fishes via precise targeting of the causative genes that are relevant to economically important traits.

## Materials and Methods

### K-mer analysis for estimating the genome size

We generated a total of 426,220,510 sequence reads for one female individual corresponding to 63.92 Gb of clean data from two 270-bp short insert-size libraries. To estimate the genome characteristics, we obtained 52,515,035,178 21-mers using these short reads. The genome size (G) is correlated with the 21-mer number (N) and the peak of 21-mer frequency (D). Their relationship can be expressed in an empirical formula: G = N / D. The kmer density distribution showed that the peak emerged at about 80 kmer-coverage. The estimated genome size was 656.46 Mb (Figure S1), which was close to the assembled genome size. By taking the estimated genome size as a reference, total sequence data accounted for an ~247-fold genome coverage (Table S1a).

### Genomic DNA preparation and sequencing

One healthy female individual collected from Nanao Town, Dapeng District, Shenzhen City, Guangdong Province, China was used for *de novo* genome sequencing. Genomic DNA was extracted from the muscles of the female adult golden pompano using the Blood & Cell Culture DNA Mini Kit (Qiagen). DNA concentrations and quality were measured using a NanoDrop 2000 (Thermo) and a Qbit Fluorometer (Thermo Fisher), respectively. Library preparation and quality assessments were conducted according to the standard protocol of the HiSeq 4000 platform (Illumina, USA). Three short fragment paired-end libraries (mean insert sizes: 270 bp and 500 bp) and nine mate-paired libraries (mean insert sizes: 3 Kb, 4 Kb, 8 Kb, 10 Kb, 15 Kb, and 17 Kb) were constructed using the Illumina standard pipeline (Table S1a). All of the libraries were sequenced on the Illumina HiSeq 4000 platform (Illumina, USA) to generate approximately 160 Gb data representing 247.83-fold genome coverage (Table S1a).

### Genome assembly

Raw data were filtered by the following criteria: (1) paired-reads with an ambiguous nucleotide ratio greater than 5% were deleted; (2) low-quality reads with a mean PHRED score < 20% (referring to sequencing error rates < 1%) were filtered; (3) reads with adapter contamination were deleted; (4) duplicate reads for mate-end libraries were deleted. We generated approximately 160 Gb of clean data, accounting for approximately a 246-fold coverage of the estimated genome size (Table S1a). Illumina sequences from short insert fragments and mate-paired reads including 270 bp, 500 bp, 3 Kb, 4 Kb, 8 Kb, 10 Kb, 15 Kb, and 17 Kb insert size libraries were initially assembled with ALLpaths-LG (allpathslg-52488)^53^ and then scaffolded using SSPACE-standard (v3.0) ^54^ (Table S2a). Taking advantage of single-molecule real time (SMRT) sequencing technology, ~4.32 Gb PacBio RSII long reads were generated from the same individual for gap filling to further improve the genome contiguity (Tables S1b and S2a).

### Genomic map construction using Bionano data

To develop a robust physical map of the golden pompano genome that could help place and order contigs/scaffolds on chromosomes and to determine the physical lengths of gaps between the contigs, we constructed BioNano optical genome map libraries for the sequenced individual. The enzyme density and distribution assessment of genome sequences were estimated using Label Density Calculator v1.3.0 (BioNano Genomics, USA). High-molecular weight DNA was extracted from blood with a Bionano Plant Tissue DNA isolation kit (Bionano Genomics) and digested with Nt.BspQI nickase (New England Biolabs). After labeling and staining, DNA was loaded onto the Irys chip for sequencing. The basic processing of BioNano raw data was conducted using the IrysView v2.5.1 package (BioNano Genomics). Molecules whose length was above 100 kb (with label SNR ≥ 3.0 and average molecule intensity < 0.6) were retained for further genome map assembly. A total of 130 Gb high quality optical molecules (508,213 molecules) that accounted for a 200-fold genome coverage (Table S1c) were collected and converted into a BNX file by AutoDetect software to obtain basic labeling and DNA length information. The high-quality DNA molecules in BNX format were aligned, clustered, and assembled into a BNG map by using the Bionano Genomics assembly pipeline. The alignment of sequence assemblies with the BNG map were computed with RefAligner, and visualization of alignment was performed with snapshot in IrysView. Based on the Label position on the single DNA molecules, *de novo* assembly was performed by a pairwise comparison of all single molecules and overlap-layout-consensus path building, which was performed by Irys-scaffolding in the IrysView v2.5.1 package. We considered only molecules containing more than seven nicking enzyme sites for assembly (min label per molecule: 8). A *P* value threshold of 1e-8 was set during the pairwise assembly, and values of 1e-9 for extension and refinement steps and 1e-12 for merging contigs were adopted. The resulting physical map covered approximately 725.16 Mb. We generated 580 optical maps with N50 of 2.05 Mb (Table S1c). The high-quality optical map was used for genome curation and hybrid assembly with SMRT-based assembly, combining the meta-pair (MP) links and HiC data.

### Hybrid assembly and gap polishing

Combination of the genome maps with the initial assembly to produce a hybrid scaffold was performed sequentially. From the comparison between the contigs/scaffolds and optical maps by RefAligner v5122 (Bionano genomics software downloads: https://bionanogenomics.com/support/software-downloads/), we assembled the 1,490 hybrid scaffolds based on a genome map hybrid assembly with an N50 size of 21.02 Mb (Table S2a). The intra-scaffold gaps were further filled with corrected PacBio long reads using PBjelly (PBSuite v15.2.20. beta)^55^ (Figure S2 and Table S2a).

### Chromosome construction using Hi-C links

Hi-C technology enables the generation of genome-wide 3D proximity maps and is an efficient and low-cost strategy for clustering, ordering, and orienting sequences during pseudomolecule construction^56^. We constructed Hi-C fragment libraries from muscle tissue for the same female individual. Libraries (based on HindIII) of fragments ranging from 300 to 700 bp sizes were constructed and sequenced using the Illumina X-TEN platform (Illumina, USA). Mapping of Hi-C reads and assignment to restriction fragments were performed as described elsewhere^56^. Briefly, adapter sequences of raw reads were trimmed with cutadapt v1.0, and low-quality paired-end (PE) reads were removed for raw data processing. The 24.98 Gb clean Hi-C reads, accounting for ~38.7-fold coverage of the golden pompano genome, were mapped to assembly results using bwa align v0.7.10^57^ with default parameters (Table S1d). Only uniquely aligned pairs of reads whose map quality was > 20 were considered for chromosome construction. Duplicate removal, sorting, and quality assessment were performed with HiC-Pro v2.8.1^58^ with the command “HiC-Pro_2.10.0/scripts/mapped_2hic_fragments.py -v -S -s 100 -l 1000 -a -f -r -o”. After data assessment, the percentage of valid interaction pairs of Hi-C data was 89.8%. Raw counts of Hi-C links were aggregated in 50-kb bins and normalized separately for intra- and inter-chromosomal contacts using HiC-Pro^58^. A pre-assembly was performed for error correction (chimeric errors) from contigs/scaffolds. Any two bins showing an inconsistent connection with information from the raw sequences were split into two fragments for reassembling. These corrected contigs/scaffolds were then assembled using LACHESIS^56^. Hi-C data were mapped to chromosomes, and placement and orientation errors exhibiting obvious discrete chromatin interaction patterns were manually adjusted. We clustered the sequences into an initial set of 24 groups according to thresholds of the contact frequency. Then, LACHESIS was used to assign the order and orientation of each group. The vast majority (99.56%, 636.6 Mb) of the assembled sequences were anchored onto the 24 pseudo-chromosomes (Figure S3 and Table S2b).

### Quality assessment for assembly results

The genome completeness and accuracy were assessed by four data sets: Pacbio sub-reads, BUSCO^59^, RNAseq data and Hi-C data. The assembled contigs were supported by more than 98% average sequence identity and nearly 100% coverage of the top 10 sub-reads (with sequence length > 30 kb) using BLASR version 1.3.1^60^ (Table S3a). The BUSCO v3.0.2 program^59^ was run against the Actinopterygii dataset (4,454 conserved protein models) (https://busco.ezlab.org/frame_wget.html) with the default parameters, resulting in 97.16% complete BUSCOs (Table S3b). For gene region completeness assessment, more than 99.46% of the unigenes *de novo* assembled from the transcriptome of intestines, middle kidney, head kidney, muscle, heart, and eggs had best hits on single contigs (Tables S3c and S3d). To assess the quality of the assembly, the interaction matrix of each chromosome was visualized with a heatmap at a 100 kb resolution.

### Repeat sequence analyses

A combination of *ab initio* and homology-based strategies was used for repeat content annotation in the golden pompano genome. The *ab initio* repeat annotation was carried out by successively using RepeatModeler (version1.0.5) (http://www.repeatmasker.org/RepeatModeler/) and RepeatMasker (version 3.3.0). The golden pompano repeat library was constructed by RepeatModeler, including two complementary programs, RECON and RepeatScout^61^. LTR-FINDER^62^, MITE-hunter^63^, and PILER-DF ^64^ were also used for *ab initio* prediction. To obtain the intact LTR consensus sequences, we integrated the LTR-FINDER results and removed false positives from the initial predictions by the LTR_retriever pipeline^65^. The yielded consensus sequences were manually checked by aligning to the Repbase transposable element library (version 16.0) The pompano repeat library consisted of 1,252 consensus sequences and their classification information, which was used to run RepeatMasker on the assembled scaffolds. The combination of results from different *ab initio* tools was used to construct a new repetitive sequence database. This database was then merged with the Repbase known transposable element library (version 16.0)^66^ and classified into different categories by the PASTEClassifier.py^67^script included in REPET v2.5^68^. The pompano repeat library was used to run RepeatMasker^69^ on the golden pompano genome. Approximately 22.46% repeat elements (~145.11 Mb sequence length) were annotated for the golden pompano (Table S4).

### Identification of non-coding RNA genes

Different types of non-coding RNA in the golden pompano genome were identified and classified to family and subfamily. The tRNAscan-SE (version 1.23)^70^ was applied to detect reliable tRNAs in the golden pompano genome. The miRNAs were identified by homology searches taking the miRBase (Release 21)^71^ as a reference, with one mismatch allowed. Then, secondary structures of the putative sequences were predicted by miRDeep2^72^. Finally, putative miRNAs with hairpin structure were considered as confident. We adopted the homology strategy to inspect other types of non-coding RNA by comparing the secondary structure similarity using Infernal (e value ≤ 0.01)^73^ based on the Rfam database (release 12.0)^74^. In total, six types of ncRNA were identified, namely, tRNAs: transfer RNAs, rRNA: ribosome RNA, miRNAs: microRNAs, sRNA: small RNA, snRNAs: small nuclear and small nucleolar RNAs, and lncRNA: long non-coding RNA. Meanwhile, 1,274 ncRNAs with a total length of 128,846 bp for the female were identified in the golden pompano genome (Table S5)

### Protein coding genes annotation

A combination of homology-based, *de novo* gene prediction and RNA-seq data strategies was adopted for gene prediction. All of the predicted gene structures were integrated into weighted consensus gene structures using EVidenceModeler (EVM, version 1.1.1)^75^. Teleost proteins of three high-quality annotation collections were obtained from the NCBI database (*Larimichthys crocea:* assembly L_crocea_2.0, ASM74293v1; *Danio rerio:* Zv9 assembly, GCA_000002035.2; *Notothenia coriiceps:* assembly NC01) and used to perform homology-based prediction. Then, we used GeMoMa (version 1.3.1)^76^ to predict the corresponding gene structure. For *de novo* prediction, we used Augustus (version 2.4) with parameters trained by unigenes from transcriptome data of golden pompano pooled tissues. Regarding the third approach, unigenes were aligned to the genome assembly using BLAST (identity ≥ 0.95, coverage ? 0.90) and then filtered using PASA v2.0.4 (http://pasapipeline.github.io/). Taking into account the weights of the three methods, all of the predicted gene structures were integrated into consensus gene structures using EVidenceModeler (EVM)^75^. We also mapped pooled transcriptome data to the reference genome using TopHat^77^and assembled transcripts with Cufflinks^77^. Transdecoder^78^was then applied to identify the structures of new gene models and new transcripts derived from Cufflinks. To obtain high-confidence gene models, the gene set was filtered by the following steps: 1) CDS lengths shorter than 300bp were removed; 2) the CDS whose lengths were not triplets were deleted; and 3) gene models with stop codons occurring in the CDS region were filtered. Finally, 24,186 protein-coding genes of the female genome were annotated for the golden pompano, a number that is comparable to the 25,497 genes of the zebrafish (Table S6a). Gene models were annotated by homology alignments against several databases using BLASTP from the BLAST+ package20 (E-value = 1e-5)^79^, including nr, nt, Swissprot, and TrEMBL. InterProScan (v4.3) was used to collect domain information and GO terms annotation for the gene models. Meanwhile, KAAS (KEGG Automatic Annotation Server) was used for the KEGG pathway annotation. In total, 22,243 genes for the male and female could be assigned to a specific function, accounting for 97.48% of the male and female whole gene set (Table S6b).

### Pseudogenes prediction

We carried out a whole-genome search to identify pseudogenes in the golden pompano. Only candidate pseudogenes with frameshift and/or premature stop codon occurrence were considered in the study. Proteins of zebrafish, large yellow croaker, and golden pompano were aligned to the golden pompano reference genome using GenBlastA (version 1.0.4)^80^ for candidate homology region identification. The candidate pseudogenes were distinguished via genewise v2.4.1^81^ with frameshift and/or premature stop codon occurrence in the coding region. After redundant filtering and manual inspection, a total of 151 confident pseudogenes were found for the golden pompano (Table S5).

### Transcription factor (TF) identification

The TFs in *T. ovatus* and other fishes were predicted using two methods, homology-based prediction and conserved domain identification. Briefly, GeMoMa v1.3.1^76^ was used to perform homology-based prediction with the TFs of zebrafish as a reference. The sequence similarity of TFs between zebrafish and *T. ovatus* was analyzed using BLASTP with parameter e-value =1e-5. The homologs with query coverage < 0.8 or target coverage < 0.8 were removed. Next, all of the predicted genes from the genome were used to annotate the domain using the hmmscan module of hmmer v3.0^82^ against the database PFAM v 27.0^83^. Genes with conserved domains of zebrafish TFs were retained as candidate TFs. Lastly, we combined the result of the above-mentioned TF identification method to generate 1,364 TFs within 45 families on the female genome (Table S17).

### WGD events

The all-to-all BLASTP program was used to identify homologous pairs, and syntenic blocks were recognized using MCscanX^84^ with parameters E_VALUE=1e-05, MAX GAPS=25. and MATCH_SIZE=5 in *S. salar*, *T. ovatus*, *M. miiuy*, *O. mossambicus*, *S. salar*, *D. rerio*, and *L. oculatus*. Synonymous substitution rates (Ks) of ohnolog pairs were calculated using codeml in the PAML package v4.7b^85^. The Ks peaks of ohnologs represent recent and ancient WGDs, and the Ks peaks of orthologs indicate speciation events. Ks ≤ 0.05 of gene pairs within *S. salar* were assigned to the Salmonid-specific WGD (Ss4R) event, while Ks > 0.05 were assigned to the teleost specific WGD (Ts3R) event. The mutation rate was 1.5e-8 substitutions per site per year. The formula t = peak Ks/2r was used to estimate the occurrence time of whole-genome duplication events, where t is time, Ks is the peak value of the substitution rate, and r is the neutral substitution rate (r = 1.5e-8).

### Phylogenetic tree construction and diverge time estimation

The longest transcript was chosen to represent the gene. To define gene families that descended from a single gene in the most recent common ancestor, OrthoMCL (version 2.0.9; mcl inflation factor 1.5)^86^ methodology was used to cluster gene families. We downloaded the protein-coding sequences of *Danio rerio* (Zv9 assembly, GCA_000002035.2), *Larimichthys crocea* (assembly L_crocea_2.0), *Lepisosteus oculatus* (GCA_000242695.1), *Miichthys miiuy* (GCA_001593715.1), Nile Tilapia (GCF_001858045.2), *Notothenia coriicep* (GCA_000735185.1 NC01), *Oncorhynchus mykiss* (GCF_002163495.1), *Salmo salar* (GCF_000233375.1), *Thunnus orientalis* (GCA_000418415.1), and *Seriola dumerili* (GCF_002260705.1) from the Ensemble database (release 56) and NCBI. A total of 1,447 single-copy orthologs shared by 11 species were obtained through gene family clustering using OrthoMCL v2.0.9. The protein sequences of single-copy orthologs were aligned by MUSCLE v3.8.31^87^ and concatenated into a super-gene sequence. We then constructed a phylogenetic tree using the maximum likelihood (ML) algorithm with the JTT amino acid substitution model implemented in phyML software^88^. The divergence time was estimated using the MCMCtree program in the PAML v4.7b (Phylogenetic Analysis of ML) package. Five calibration points (*S. salar* vs. *O. mykiss:* 28.10-28.40 MYA, *T. ovatus* vs. *S. dumerili:* 57-63 MYA, *M. miiuy* vs. *L. crocea:* 81-101 MYA, *T. orientalis* vs. *S. dumerili:* 117-128 MYA, and *L. oculatus* vs. *S. dumerili:* 305-360 MYA) derived from the TimeTree database (http://www.timetree.org/) were applied to constrain the divergence times of the nodes.

### Gene family expansion/contraction analysis

We applied the likelihood model implemented in the CAFE v2.2^89^package to identify the expanded and contracted gene families along each branch of the phylogenetic tree. The topology and branch lengths of the phylogenetic tree were taken to infer the significance of changes in gene family size. The significant levels of expansion and contraction were set at 0.05.

### Fish materials and F_1_ mapping population construction

Both parents used in this study were selected from 26 healthy and non-deficit golden pompano at sexual maturity (5–7 years old, 8–13 kg weight) from Nanao Town, Dapeng District, Shenzhen City, Guangdong Province, China. After intensive cultivation, all these golden pompanos were used as broodstocks for family construction. Before marking and spawning induction, the broodstocks were exercised twice a day for 2–3 days to increase their abilities to adapt to environmental changes and reduce subsequent operation stress responses. Brightly colored and easily distinguishable ring tags were chosen and fixed on the tail stalk of the golden pompano. HCG (500–1000 U/kg) and LRH-A2 or LRH-A3 (2–7μg/kg) were injected into the dorsal muscle or caudal stalk muscle of the broodstocks. When the males were chasing the females, the colored ring logs in the tails of the females were recorded. The tail-chasing broodstock pair were carefully picked up and put into a separate cage or pool. Some immature golden pompano were put into the cage or pool to form a pair-mating environment. Twelve to twenty-four hours after the spawning, the floating fertilized eggs of each broodstock pair were carefully washed and removed into an incubation bucket (22–32°C) with sufficient oxygen (4 mg/L) until the fertilized eggs hatched (intensity less than 0.6 million eggs per m^3^). All of the hatched offspring were cultured in marine (salinity 25–35‰) aquaculture ponds as an F_1_ hybrid population (> 2000) of each broodstock pair. Samples from pairs of parents with the largest size/weight difference were selected to perform the genetic map construction and QTL mapping. The appearance data of the F_1_ population are shown in Fig 3A. The rearing of larvae and adults was performed according to standard protocols.

### Phenotype recording

Two broodstock pairs were successfully mated, and one of their offspring was selected to carry out the subsequent experiments. A total of 200 six-month-old progeny from this F_1_ population were randomly selected and numbered, and their total length (TL), body length (BL), body height (BH), and body width (BW) at three time points one month apart were separately measured and recorded. The dorsal muscle or caudal stalk muscle of each progeny was harvested and stored at −80°C. Phenotypic data for all growth traits were analyzed using GraphPad Prism 7.0 (GraphPad Software, San Diego, CA) software. Statistical evaluation was carried out by using Student’s t tests, ANOVA, or *χ*^2^ tests as appropriate. Statistical significance was set at *p* < 0.05.

### DNA extraction and whole-genome resequencing

Dorsal muscle from the parents and their 200 progeny (from Nanao Town, Dapeng District, Shenzhen City, Guangdong Province, China) and 147 three-to five-month-old fish from a natural population (from a clean and pollution-free offshore marine aquaculture farm in Bailong Town, Fangcheng City, Guangxi Province, China) were used to extract genomic DNA. The DNA was extracted using the Blood & Cell Culture DNA Mini Kit (Qiagen) kit according to the manufacturer’s protocol. The concentration and quality of the total genomic DNA were determined by using 1% agarose gel electrophoresis and an ND-1000 Spectrophotometer (NanoDrop, USA). DNA samples were dissolved in ddH2O and stored at −80°C until use. Illumina sequencing DNA libraries were constructed according to the manufacturer’s specifications (Illumina, CA, USA). Briefly, Covaris S220 (Covaris, MA, USA) was used to shear the genomic DNA into approximately 350 bp fragments. Then, fragments were end repaired with an additional base A and sequencing adapter. Finally, target fragment selection and PCR enrichment were performed, and clustering was generated with Cbot. An Illumina HiSeq Xten platform (Illumina, San Diego, CA, USA) was used to perform paired-end 100 bp sequencing in two lanes. Raw reads were filtered as follows: the adapter sequence was removed, and low-quality reads (the number of bases with a quality value Q ≤ 10 accounting for more than 50% of the entire read) were deleted. If the proportion of N (the specific base type could not be determined) on a read was greater than 10%, the paired-end reads were filtered out. Finally, 8.3Gb (female parent), 8.6Gb (male parent), and 203.38Gb (200 offspring) of high-quality clean data were obtained for subsequent analyses.

### SNP calling and genotyping

Raw data obtained by Illumina sequencing were filtered to obtain clean data. The data filtering criteria were as follows: the adapter sequences were removed, and the paired-end reads were deleted if the proportion of N on a read (the specific base type could be determined) was greater than 10%. Low-quality reads (the number of bases with a quality value Q ≤ 10 accounting for more than 50% of the entire read) were removed. Then, the clean reads were mapped onto the golden pompano reference genome, which was sequenced by our research group (deposited at DDBJ/ENA/GenBank under the accession WOFJ00000000 under BioProject PRJNA574895) using the Burrows-Wheeler Alignment tool^57^. The duplicated reads caused by PCR amplification were eliminated by the Mark Duplicate function of the Picard program (http://sourceforge.net/projects/picard/). The Genome Analysis Toolkit software (GATK)^90^ was used to call SNPs and insertions and deletions (InDels) between the reference genome and all sequenced samples._In order to ensure a high quality SNP dataset, SNPs were filtered as follows: two SNPs with 5bp were discarded; a SNP within 5bp near an InDel was filtered out;_two InDels at distances less than 10bp were discarded^91^. The Circos^15^ program was used to present the SNP and InDel data annotated with SnpEff^92^. SNPs screening between parents and genotyping of F_1_ offspring were carried out based on the above-obtained SNP dataset. The SNP screening was performed as follows: SNPs with less than four-fold depth were deleted; SNPs on unanchored contigs or scaffolds were discarded; the homozygous SNPs in parents were also discarded. Genotyping of the F_1_ population was performed according to the parental SNPs. The filtered SNP markers were divided into linkage groups (LGs) based on their locations in the reference genome, and the linkage relationships between markers on each chromosome were tested by two-points analysis. The missing genotyping imputation and the genotyping errors were corrected by SMOOTH^15, 93^.

### Genetic linkage map construction

After completing the mark filling and correction, Bin division was carried out according to the recombination and exchange of the offspring. All of the genotypes of F_1_ individuals were arranged according to the physical positions on the chromosomes. A genotyping transition in any sample was considered as a recombination breakpoint. The SNP between the recombination breakpoints was classified as a bin^94^, and non-recombination events could be observed in a bin. The bin was used as a mapping marker to construct a genetic map. A final group of bin markers was applied to construct a linkage map using HighMap software^95^. Genetic distances were calculated using the Kosambi mapping function^96^. The heatmap of adjacent markers and the degree of collinearity between the genetic map and the reference genome (Spearman correlation coefficients) were used to evaluate the quality of the genetic linkage map.

### QTL mapping and fine positioning

Combining the golden pompano phenotypic data and high-density genetic linkage map, the interval mapping (IM) in R/qtl was used for QTL mapping using a threshold LOD score of 3.0^97^. In order to verify the QTL loci, 10 KASP markers were explored based on the SNPs within the QTL loci. Genotyping was carried out in 64 individuals with evenly distributed extreme phenotypes in the F_1_ population, which had no shared samples with the QTL mapping population, and the correlation between markers and phenotypes was confirmed by using Wilcoxon tests. In order to finely locate the main effect QTLs, we developed 14 KASP markers in the bin marker of the above-confirmed QTL. A total of 147 individuals in a natural population were randomly selected and genotyped, and their total length (TL), body length (BL), body height (BH), and body width (BW) at two time points with two months apart were separately measured and recorded. After genotyping of the147 natural population individuals, the correlations between the SNP HapQTL^98^ and the phenotypes were measured by using target-fragment sanger sequencing and the T-test method. The mapping interval was determined according to the SNP position of the finely positioned SNP marker among the bin markers on the reference genome. Non-synonymous mutations and mutations in the promoter region of the gene (within 2Kb upstream of the start codon) in the mapping interval were regarded as potential casual mutations. The related genes in the mapping interval with potential casual mutations were then annotated by blasting the Gene Ontology (GO)^99^ and Kyoto Encyclopedia of Genes and Genomes (KEGG)^100^ databases. The genes related to growth in the annotation results were considered as candidate genes that need to be verified. Candidate genes are listed in Table S23. SNP validation of the candidate gene gpsstr1 was performed by sanger sequencing. Simply, muscle samples were collected from 64 F_1_ population and 147 natural populations. The gDNA was extracted by the identical method in “DNA extraction and whole-genome resequencing” section. Then the PCR was performed to amplify the 2000 bp length fragment of the promotor region of gpsstr1 gene. PCR products were purified and sequenced by sanger sequencing method. (See Table S24 for PCR primers for sanger sequencing).

### Quantification of candidate genes in the F_1_ and natural populations

The cDNA of the candidate gene gpsstr1 was quantified by qPCR amplifications.. Briefly, muscle samples were collected from 200 F_1_ progeny and 147 individuals from the natural population. Total RNAs were then extracted from muscles in Trizol (Roche) reagent according to the manufacturer’s instructions. RNAs were reverse transcribed by using the PrimeScript RT reagent Kit with gDNA Eraser (TAKARA, Kyoto, Japan). Quantification of candidate genes was conducted in triplicate by using Bio-rad iQ SYBR Green Supermix (Bio-rad, Hercules, CA, USA). Relative gene quantification was calculated according to the comparative threshold cycle method (2^-ΔΔCt^) with the 18S gene as a control. (see Table S26 for gpsstr1 primers).

### Zebrafish maintenance and microinjection

All of the wild-type AB strain zebrafish were raised at 28-28.5°C on a 14 h light/10 h dark cycle at Shanghai Model Organisms Center, Inc, which is accredited by the Association for Assessment and Accreditation of Laboratory Animal Care (AAALAC) International. After natural mating, the obtained embryos were maintained at 28.5°C in 0.2% deionized instant ocean salt water and staged based on previous research^101^. All morpholinos (MOs) were designed and synthesized by Gene Tools, LLC (http://www.gene-tools.com/). Antisense MOs were microinjected into fertilized one-cell stage embryos according to the standard protocols^102^. For *sstr1a* knockdown, the sequence of the *sstr1a* translation-blocking was 5’-AAGGTGTCGTTGGGCAGCATTCC −3’ (ATG-MO). The sequence for the standard control MO was 5’-CCTCTTACCTCAGTTACAATTTATA −3’ (Gene Tools). The amounts of the MOs used for microinjection were as follows: Control-MO and ATG-MO, 4 ng per embryo. The CDS region of *sstr1a* cDNA, including the *sstr1a*-ATG-MO target sequence, was cloned in the frame into pcDNA3.1-EGFP for testing the effectiveness of *sstr1a* MOs. For *gsk3β* knockdown, the sequence of the *gsk3ba* splice-blocking morpholino was 5’-CTGTCTCGGTCTTACCTTAAATCGC −3’ (E2I2-MO). The sequence for the standard control morpholino was 5’ - CCTCTTACCTCAGTTACAATTTATA −3’ (Gene Tools). The amount of the MOs used for injection was as follows: Control-MO and E2I2-MO, 4 ng per embryo. Primers spanning *gsk3ba* exon 1 (forward primer: 5’-TTCGGCAGCATGAAAGTC −3’) and exon 4 (reverse primer: 5’-TAGTGTCTTGCCACTCTGTA −3’) were used for RT-PCR analysis for confirmation of the efficacy of the E2I2-MO. The primer *ef1α* sequences used as the internal control were 5’-GGAAATTCGAGACCAGCAAATAC −3’ (forward) and 5’-GATACCAGCCTCAAACTCACC −3’ (reverse). At 2-dpf, embryos were dechorionated and anesthetized with 0.016% MS-222 (tricaine methane sulfonate, Sigma-Aldrich, St. Louis, MO). Zebrafish were then oriented on lateral side (anterior, left; posterior, right; dorsal, top) or dorsal side, and mounted with 3% methylcellulose in a depression slide for observation by light or fluorescence microscopy. The phenotypes of body lengths, tail, and eyes were analyzed.

### Gene silencing

The siRNA probes were designed and synthesized according to the human sstr1 gene (NM_001049.3). The probes are listed in Table S26. The 293T cells (ATCC, Manassas, Virginia, USA) were cultured in DMEM (Hyclone, USA) with 5% FBS (Gibco BRL. Co. Ltd.) and 1% penicillin-streptomycin (Sangon Biotech, China.) at 37°C in a 5% CO2 incubator. Three experimental groups, 293T, 293T+si-sstr1 NC, and 293T +si-sstr1 were set. Thirty picomoles of si-sstr1 per well were transferred into 293T cells in 24-well plates (Corning-Costa) by using 9 μl Lipofectamine RNAi MAX Reagent (Invitrogen, USA). Cells were centrifuged and collected for protein extraction after 12-24h incubation.

### RNA-seq

Zebrafish embryos injected with *sstr1a* MO and control MO at 2 dpf were collected for RNA-seq analysis. Total RNAs were extracted and purified using an RNAqueous Total RNA isolation kit (Cat. No. AM1912, Thermo Fisher). DNA contamination was eliminated by DNase digestion. The quality and quantity of RNA were determined by using an Agilent 2100 BioAnalyzer (Agilent Technologies, Santa Clara, CA). Transcriptome libraries were prepared with a TruSeq RNA library Prep kit v2 (Cat. No. RS-122-2001, Illumina) according to the manufacturer’s protocol and sequenced at the CCHMC Core Facility using Illumina HiSeq 2500 sequencing system (Illumina) to generate 100 bp paired-end reads. Fastqc [http://www.bioinformatics.babraham.ac.uk/projects/fastqc/] and trimmomatic [http://www.usadellab.org/cms/?page=trimmomatic] were applied to check the quality of the reads and to filter the low-quality reads. The clean reads were mapped to the latest Zebrafish genome assembly GRCz10 at default thresholds by using RSEM [http://deweylab.github.io/RSEM/]. TopHat v2.0.9 and Cufflinks were used to identify the mRNA levels, which were normalized by the Fragments Per Kilobase of exon model per Million mapped reads (FPKM). Differential expression of genes was analyzed by using CSBB’s [https://github.com/skygenomics/CSBB-v1.0]. ToppGene [https://topgene.cchmc.org/] was used to perform GO annotation and KEGG pathway annotation.

### Quantitative real-time PCR

Total RNA was extracted from golden pompano tissues or embryos or zebrafish embryos in Trizol (Roche) according to the manufacturer’s instructions. RNAs were reverse transcribed by using the PrimeScript RT reagent Kit with gDNA Eraser (Takara). Expression of each gene was examined in triplicate by using Bio-rad iQ SYBR Green Supermix (Bio-rad) with detection on the Realplex system (Eppendorf). Relative gene quantification was based on the comparative threshold cycle method (2^-ΔΔCt^) by using the *ef1α* gene as a control. All of the primers are listed in Table S26.

### Western blotting assays

Muscles from fast/slow growing golden pompano, zebrafish tissues from *sstr1a* knockdown and control group, or 293T cells in the sstr1 silenced group and control were treated with 1 mL of tissue lysate (150 mmol/L NaCl, 50 mmol/L Tris, 0.1% SDS, 5 mmol/L EDTA, 5 μg/mL aprotinin, 1% NP-40, and 2 mmoL/L PMSF followed by lysis with protein lysate at 4°C for 30 min). All of the samples were centrifuged at 12,000 r/min at 4°C for 30 min, and the supernatant was collected to detect the protein concentration by using a BCA kit (CWBIO. Co., Ltd., Shanghai, China). All of the samples were resolved by SDS-PAGE using a NuPAGE 4–12% gel (Life Technologies). All of the proteins were transferred onto a nitrocellulose filter (BioRad, Hercules, CA, USA) and sealed by 5% dried skimmed milk at 4°C overnight. The membranes were incubated with diluted primary rabbit polyclonal to Somatostatin Receptor 1/SSTR1 (ab140945, Abcam, UK) (1:1000), IGF1(DF8564, Affinity, Biosciences. OH. USA) (1:1000), DKK1 (ab61275, Abcam, UK) (1:1000), GSK3 (alpha+beta) (ab68476, Abcam, UK) (1:1000), WNT9A (PA5-47464, Invitrogen, USA) (1:1000), β-Catenin (GTX61089, Genetex, USA) (1:1000), AXIN2 (DF6978, Affinity, Biosciences. OH. USA) (1:1000), and GAPDH (ab8245, Abcam, UK) (1:1000) antibodies overnight at 4°C. The membranes were treated with IgG-HRP secondary antibody (1: 2000, CWBiotech., Ltd., Beijing, China) and incubated at 37°C for 2 h. The membrane was soaked in an enhanced chemiluminescence (ECL) kit (CW Biotech., Ltd., Beijing, China) according to the manufacturer’s instructions.

### Statistical analysis

All data are presented as mean ± SEM. Statistical analysis and graphical representation of the data were performed using GraphPad Prism 7.0 (GraphPad Software, San Diego, CA). Statistical evaluation was performed by using a Student’s t test, ANOVA, or *χ*^2^ test as appropriate. *p* value of less than 0.05 was considered statistically significant. Statistical significance is indicated by * or *p* value. * represents *p* < 0.05, **represents *p* < 0.01, and *** indicates *p* < 0.0001. The results are representative of at least three independent experiments.

## Acknowledgements

We thank Mr Zhuanbin Wu for zebrafish analysis advices and technical consults.

## Author contributions

H.L.L and X.H.C designed the scientific objectives and oversaw the project. H.L.L., X.H.C., Y.Z.Z., J.X.P., and Y.L discussed the primary ideas of the article. Y.D.Z, J.X.P., Y.L., Y.H., Q.Y.L., P.P.H., C.L.Y., P.Y.W., X.L.C., and P.F.F. collected samples for sequencing DNA and RNA. C.M.J. and their colleagues performed genome sequencing, assembly and annotation. C.M.J. and H.Y.Y performed phylogenomic and whole genome duplication evolution analysis. Y.H.X, H.L.L and Y.D.Z analyzed the QTL mapping and fine mapping and candidate screening. C.M.J. H.Y.Y., H.L.L., and Y.D.Z performed RNA-seq analysis. H.L.L and Y.D.Z performed functional assay of zebrafish *sstr1a* gene. C.M.J, H.Y.Y, H.L.L., Y.H.X., and Y.D.Z prepared the supplemental data and method. C.M.J., H.L.L and Y.H.X prepared the draft manuscript with input from all other authors. H.L.L., X.H.C., Y.L., Y.H.X and H.K.Z. discussed and revised the manuscript.

## Data availability

The authors declare that all data reported in this study are fully and freely available from the date of publication. This Whole Genome Shotgun project has been deposited at DDBJ/ENA/GenBank under the accession WOFJ00000000. The version described in this paper is version WOFJ01000000. The draft genome data (genome assembling and annotations) and re-sequencing data are available under BioProject PRJNA574895.

## Supplemental material

Supplemental Figures (Figure S1-S11)

Supplemental Tables (Table S1-S26)

